# Profiling a large HIV-1 elite neutralizer cohort reveals remarkable CD4bs bNAb for HIV-1 prevention and therapy

**DOI:** 10.1101/2025.08.27.672638

**Authors:** Lutz Gieselmann, Andrew T. DeLaitsch, Malena Rohde, Henning Gruell, Christoph Kreer, Meryem Seda Ercanoglu, Harry B. Gristick, Philipp Schommers, Elvin Ahmadov, Caelan Radford, Andrea Mazzolini, Lily Zhang, Anthony P. West, Johanna Worczinski, Anna Momot, Maren L. Reichwein, Jacqueline Knüfer, Ricarda Stumpf, Nonhlanhla N. Mkhize, Haajira Kaldine, Sinethemba Bhebhe, Sharvari Deshpande, Frederico Giovannoni, Erin Stefanutti, Fabio Benigni, Colin Havenar-Daughton, Davide Corti, Arne Kroidl, Anurag Adhikari, Aubin J. Nanfack, Georgia E. Ambada, Ralf Duerr, Lucas Maganga, Wiston William, Nyanda E. Ntinginya, Timo Wolf, Christof Geldmacher, Michael Hoelscher, Clara Lehmann, Penny L. Moore, Thierry Mora, Aleksandra M. Walczak, Peter B. Gilbert, Nicole A. Doria-Rose, Yunda Huang, Jesse D. Bloom, Michael S. Seaman, Pamela J. Bjorkman, Florian Klein

**Author notes:** These authors contributed equally to this work (Lutz Gieselmann, Andrew T. DeLaitsch).

## Abstract

Administration of HIV-1 neutralizing antibodies can suppress viremia and prevent infection *in vivo*. However, clinical use is challenged by broad envelope sequence diversity and rapid emergence of viral escape^1–9^. Here, we performed single B cell profiling of 32 top HIV-1 elite neutralizers to identify broadly neutralizing antibodies (bNAbs) with highest potency and breadth for clinical application. From 831 expressed monoclonal antibodies, we identified 04_A06, a new V_H_1-2-encoded CD4 binding site bNAb with remarkable breadth and potency against extended multiclade pseudovirus panels (GeoMean IC_50_ = 0.059 µg/ml, breadth = 98.5%, 332 virus strains). Moreover, 04_A06 was not susceptible to classic viral CD4bs escape variants and maintained full viral suppression in HIV-1-infected humanized mice. Structural analyses revealed that antiviral activity is mediated by an unusually long 11-amino acid heavy chain insertion. This insertion facilitates inter-protomer contacts and interactions with highly conserved residues on the adjacent gp120 protomer. Finally, 04_A06 demonstrated high activity against contemporaneously circulating viruses from the Antibody Mediated Prevention (AMP) trials (GeoMean IC_50_ = 0.082 µg/ml, breadth = 98.4%, 191 virus strains) and *in silico* modeling for 04_A06LS predicted HIV-1 prevention efficacy of >93%. Thus, 04_A06 will provide unique opportunities for effective treatment and prevention strategies of HIV-1 infection.

## Introduction

Administration of monoclonal antibodies (mAbs) has emerged as a pivotal strategy in the treatment and prevention of infectious diseases, offering a targeted approach to combat viral infections^10–14^. Due to their capacity to neutralize diverse HIV-1 strains and subtypes, broadly neutralizing antibodies (bNAbs) represent a promising tool for immunotherapy and prevention^15–18^. bNAbs target distinct sites of vulnerability on the HIV-1 Env trimer, including the CD4 binding site (CD4bs), glycan-dependent sites at the V3 loop base and V2 region, the gp120-gp41 subunit interface, the membrane-proximal external region, the fusion peptide, and the silent face^19–24^. Located on the outer domain of the gp120 subunit, the highly conserved CD4bs is critical for virus engagement of host cells. Given its pivotal role in the viral lifecycle, many CD4 binding site-directed bNAbs display high levels of antiviral activity, and viral evasion from these bNAbs may entail significant fitness costs^25^. As a consequence, CD4bs bNAbs are a prime target for clinical evaluation and vaccine development^26–31^.

CD4bs bNAbs can be categorized into VRC01-class (e.g., VRC01, 3BNC117, N6, N49P7, NIH45-46, and VRC07-523)^32–35^ and non-VRC01-class (e.g., CH103, 8ANC131, 1-18)^36–38^ antibodies. Members of the VRC01-class are encoded by the immunoglobulin heavy chain gene segment IGHV1-2 include a five-residue light chain complementary-determining region 3 (CDRL3), and are characterized by high levels of somatic hypermutation^19,21,39,40^. Although bNAbs have shown to (i) reduce viremia, (ii) delay viral rebound, and (iii) prevent infection with sensitive viruses, treatment and prevention trials have highlighted limitations that impede the clinical applicability of bNAbs^2–7,9,17,41–46^. These include HIV-1 Env sequence diversity and pre-existing and *de novo* viral resistance^2–8,19,47–50^. Therefore, identification of bNAbs with enhanced potency, breadth, and restricted viral escape pathways is critical for the integration of bNAbs into clinical practice.

Most bNAbs in clinical testing were isolated from a small number of HIV-1 elite neutralizers^33,34,36,37,51–55^. In these donors, bNAbs typically emerge from individual or co-existing B cell lineages during coevolution between host immunity and the evading virus. However, as both elite neutralizers and bNAb lineages within these individuals are rare^53,56–62^, employing an extensive large-scale screening approach can support the discovery of new bNAbs with promising properties for clinical application.

Combining micro-scale antibody production with direct functional testing^63^, we performed detailed single-cell profiling of the largest cohort of top HIV-1 elite neutralizers studied to date (32 individuals). As a result, we identified 04_A06, a highly broad and potent CD4bs bNAb, that emerged from one of three genetically divergent B cell clones with overlapping epitope specificity for the CD4bs. 04_A06 contains a frame work region heavy chain (FWRH) 1 11-amino acid insertion, which we show by single-particle cryo-EM analysis contacts highly conserved residues (>99%) on HIV-1 Env, providing a structural explanation for its remarkable antiviral activity. This also allows 04_A06 to overcome classic viral CD4bs escape and to achieve full suppression of viremia in HIV-1_YU2_-infected humanized mice. Finally, based on high neutralizing activity of 04_A06 against transmitted viruses obtained from two antibody-mediated prevention trials^9^ (AMP) (ClinicalTrials.gov numbers, NCT02716675 and NCT02568215), modeling predicted a prevention efficacy of 93% for an extended half-life variant, making 04_A06 a promising bNAb candidate for effective treatment and prevention of HIV-1 infection.

## Results

### Large scale profiling of HIV-1 elite neutralizers reveals new HIV-1 bNAbs

We recently established an international cohort of 2,354 people living with HIV-1 (PLWH) and ranked each individual based on their serum IgG neutralizing activity against the 12-strain HIV-1 global pseudovirus panel^64,65^. From 32 elite neutralizers of our cohort, we collected large blood samples to identify new HIV-1 bNAb candidates (Fig. 1a, Supplementary Table 1). Donors were between 21 and 53 years (median of 38 years), 47% were female, and 66% were off antiretroviral therapy (ART) at the time of blood draw (Fig. 1a, Supplementary Table 1). Samples were obtained in Tanzania (44%), Germany (25%), Nepal (25%), and Cameroon (6%) (Fig. 1a, Supplementary Table 1). To isolate HIV-1 Env-reactive B cells, we performed single cell sorting using GFP-labeled BG505_SOSIP.664_ and YU2_gp140_ bait proteins^66,67^ (Extended Data Fig. 1a). HIV-1 Env-reactive IgG^+^ memory B cell frequencies ranged from 0.005–0.67% (median 0.1%) (Extended Data Fig. 1a). From a total of 5,324 isolated single B cells, we amplified 4,949 IgG heavy and 2,256 light chains (2,255 heavy and light chain pairs) by applying optimized PCR primers and protocols^63,68^. In each individual, 4.3–100% of sequences were clonally related with a mean of 18 clones per individual and a mean clone size of 5 (Extended Data Fig. 2a,b). To identify neutralizing antibodies (nAbs), we expressed 831 mAbs from 27 donors and tested each for neutralizing activity against a screening panel of six HIV-1 pseudoviruses representative of different clades (Fig. 1b,c, Supplementary Table 2)^69^. Antibodies were selected to represent all identified B cell clones, and additionally, non-clonal mAbs were included that exhibited V_H_ gene sequences with uncommon features, such as high level of somatic hypermutation (≤80% V_H_ gene germline identity), amino acid insertions or deletions, and long CDRH3 regions (≥25 amino acids in length). From all screened antibodies tested at a concentration of 2 µg/mL, 214 (25%) displayed ≥50% neutralization against at least one virus strain (Fig. 1b,c). Most nAbs demonstrated activity against a single tested strain (124/214, 58%) and only seven mAbs from only two donors (3.3%) neutralized all strains of the screening panel (Fig. 1c). Overall, enhanced potency of nAbs was associated with increased breadth (Fig. 1c). Compared to healthy reference memory B cell repertoires^70^, mAbs isolated from HIV-1 elite neutralizers demonstrated enriched V_H_ gene segments (e.g., V_H_1-69, V_H_1-2, and V_H_4-34), longer CDRH3 lengths, and higher V_H_ mutation frequencies (Fig. 1d). Within the subset of nAbs, we detected higher fractions of V_H_ gene segments V_H_5-51, V_H_1-69-2, and V_H_3-43 than in non-neutralizing mAbs (Fig. 1d). The average CDRH3 length of nAbs was slightly longer, and on average, nAbs displayed a reduced net charge but no differences in hydrophobicity when compared to non-neutralizing mAbs (Fig. 1d). In addition, nAbs acquired more V_H_ gene mutations than non-neutralizing antibodies and a higher mutation rate correlated with higher neutralizing activity (Fig. 1e, f). We conclude that anti-HIV-1 nAbs emerge from a broad set of different V genes with a preference for V_H_5-51, V_H_1-69-2, and V_H_3-43 gene segments over non nAbs and are characterized by high degrees of somatic mutations that correlate with antiviral activity^70^. Moreover, despite in-depth investigation of a large cohort of HIV-1 elite neutralizers, broad and potent anti-HIV-1 bNAbs remain exceptionally rare.

**Figure 1:**
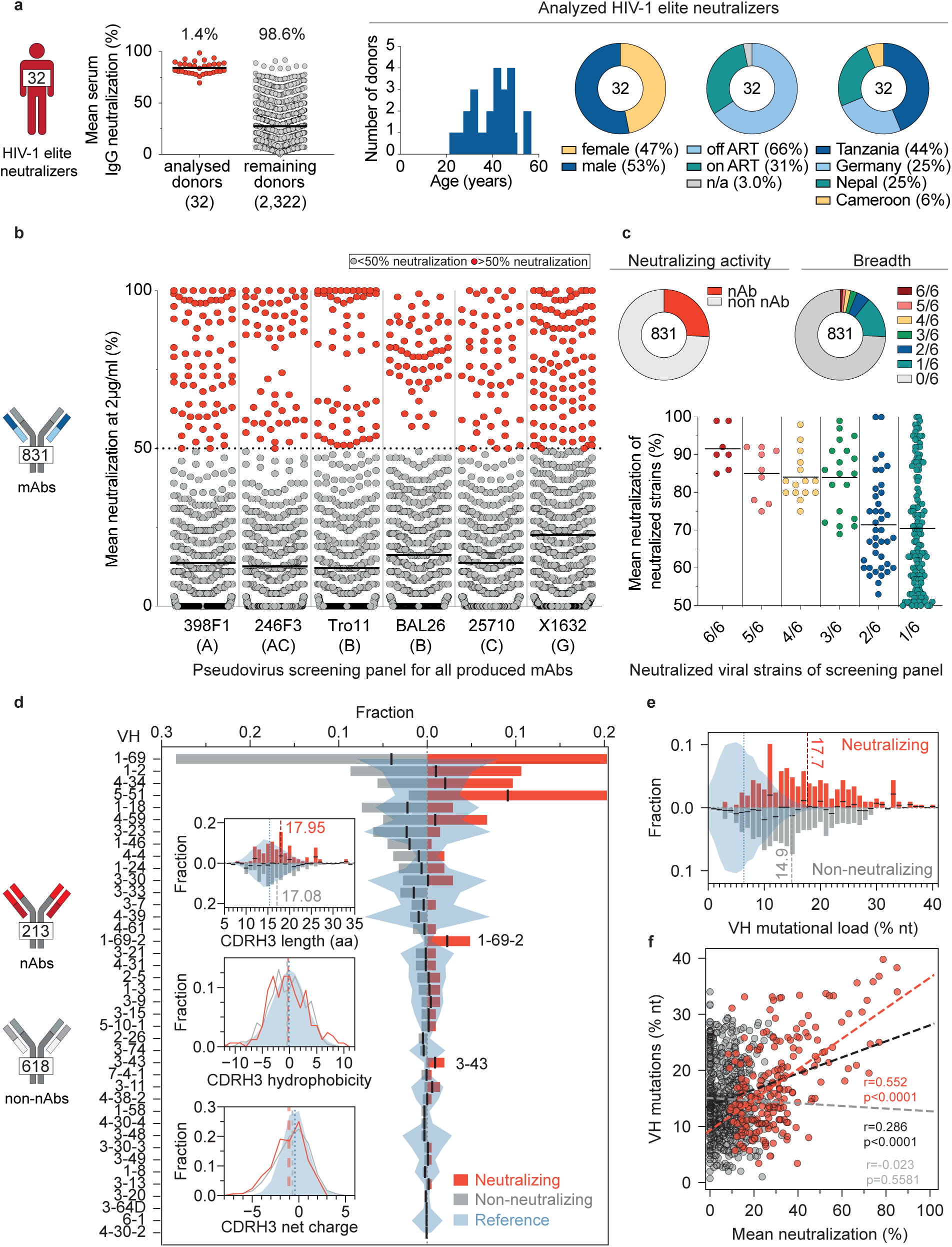
Large-scale isolation of mAbs from HIV-1 elite neutralizers. **a**, HIV-1 neutralizing serum activity and demographic characteristics of 32 HIV-1 elite neutralizers^65^. Dot plots illustrate the neutralizing activity of purified serum IgG samples of each donor against the HIV-1 global pseudovirus panel^64^. Age distribution, sex, ART status of donors and geographical origin of the samples are illustrated as pie charts and bar plots. Serum IgG samples were tested in duplicates^65^. **b**, Neutralizing activity of isolated 831 mAbs against a screening pseudovirus panel of 6 viral strains from 4 different clades. Each antibody was evaluated at a concentration of 2 μg/ml. Antibodies that achieved greater than 50% neutralization were classified as neutralizing (red; those with <50% neutralization were categorized as non-neutralizing mAbs (grey). Antibodies were tested once. **c**, Illustration of the mAb antiviral activities. Pie charts display the proportion of nAbs, their levels of breadth, and their neutralization spectrum across HIV-1 clades. Center lines in dot plots in **a**, **b**, and **c** indicate means. **d**, V_H_ gene segment usage and CDRH3 characteristics of 831 mAbs divided into nAbs (red) and non-nAbs (grey). V gene segments are ordered by the overall frequencies in all 831 antibodies. Bar graphs show fractions that were calculated for individual subsets. CDRH3 characteristics comprise lengths in amino acids, cumulative hydrophobicities (Eisenberg scale) and net charge. For comparison, NGS reference data from 48 healthy individuals^70^ (blue) is shown. **e**, V_H_ mutational load of non-nAbs (grey) and nAbs (red). **f**, Correlation between V_H_ mutations and neutralizing activity of antibodies. Pearson correlation coefficients r and two-sided p-values for nAbs (red), non-nAbs (grey) and all (black) mAbs are reported in the figure. Dashed lines show linear regressions for the respective subset. Black lines within bar graphs in **d** and **e** depict the center of combined fractions to indicate overrepresentation in either nAbs or non-nAbs. Dashed lines in **d** and **e** represent distribution means.

### Phylogenetic analyses reveal co-existence of genetically distinct CD4bs bNAbs

Among all the screened mAbs, those derived from individual EN02, a female living with HIV-1 clade C in Tanzania, exhibited the highest levels of breadth (up to 100%) and mean neutralization potency (up to 99%) against the HIV-1 pseudovirus screening panel^65^ (Supplementary Table 2). Purified EN02 serum IgG displayed broad and potent activity with a coverage of 100% and a mean serum IgG neutralization of 94% against the 12-strain global HIV-1 pseudovirus panel^64,65^ (Extended Data Fig. 3a, Supplementary Table 1). Based on serum IgG antiviral activity, EN02 was ranked as the second top elite neutralizer within our cohort of 2,354 PLWH^65^. Neutralization fingerprint analysis of purified serum IgG revealed VRC01-like activity suggesting neutralization to be predominantly mediated by CD4bs antibodies^65,71^ (Extended Data Fig. 3b).

The majority of nAbs isolated from donor EN02 were encoded by the heavy chain V gene segment V_H_1-2*07 paired with a kappa light chain derived from V_K_1-33*01, most likely containing a CDRL3 of 5 amino acids in length, a common feature of V_H_1-2-encoded CD4bs bNAbs (Fig. 2a; Supplementary Table 3). However, due to the extensive level of hypermutation the precise V_H_ gene allele assignment was ambiguous for some mAbs (e.g., sequence identities for V_H_1-2*07, *02, *04 of 61.7%, 61.4%, and 61.1% for 04_A06 determined by IMGT algorithm, respectively). The isolated antibodies were highly mutated with 60%-83% V_H_ gene germline nucleotide identities and shared only up to 49% (heavy chain) or 47% (light chain) amino acid sequence identity (Fig. 2a; Supplementary Table 3). Importantly, the isolated heavy chain sequences of nAbs varied with differences observed in the length, position, and/or sequence of amino acid insertions within the FWRH1 or FWRH3 regions, as well as in the lengths and/or sequences of the CDRH3 (Fig. 2a; Supplementary Table 3). Whereas some nAbs lacked amino acid insertions, others displayed six and four amino acid insertions in the FWRH1 and FWRH3 or an ultra-long insertion of 10 or 11 amino acids in the FWRH1 (Fig. 2a; Supplementary Table 3). These findings were indicative of parallel B cell evolution from distinct B cell progenitors. However, delineation of B cell ontogeny and clonality of the isolated nAbs was challenging due to their high mutation frequencies^72,73^ and lack of a widely accepted definition and specific criteria for identifying B cell clones. To overcome these challenges, we established the fraction of common V_H_ gene mutations of isolated bNAbs as a quantifiable and comparable metric of sequence similarity (Fig. 2b). Applying this metric, we compared mutations within bNAb sequences isolated from the same donor (intradonor similarities) as well as between bNAb sequences from different donors (interdonor similarities) encoded by the same V_H_ gene. For our analyses, we applied IGHV1-2-encoded CD4bs bNAb sequences isolated from different donors as a comparator dataset (Fig. 2b). Using interdonor similarities as a benchmark for unrelated sequences, we inferred that if two bNAbs within a donor are as dissimilar as those from different donors, they likely do not share a phylogenetic relationship and are considered part of different B cell clones that originated from distinct B cell progenitors. The analysis of intradonor similarities in sequences from donor EN02 revealed a bimodal distribution. The first peak aligned with interdonor similarities, suggesting these sequences originated from unrelated B cell clones. The second peak indicated higher sequence similarity, suggesting a clonal relationship. A threshold was defined between these peaks to differentiate clonally-related from unrelated sequences. Employing this threshold, we identified three clusters of clonally-related sequences, representing genetically-distinct B cell clones that likely arose from different B cell progenitors (Fig. 2b) but share specificity for the CD4bs.

**Figure 2:**
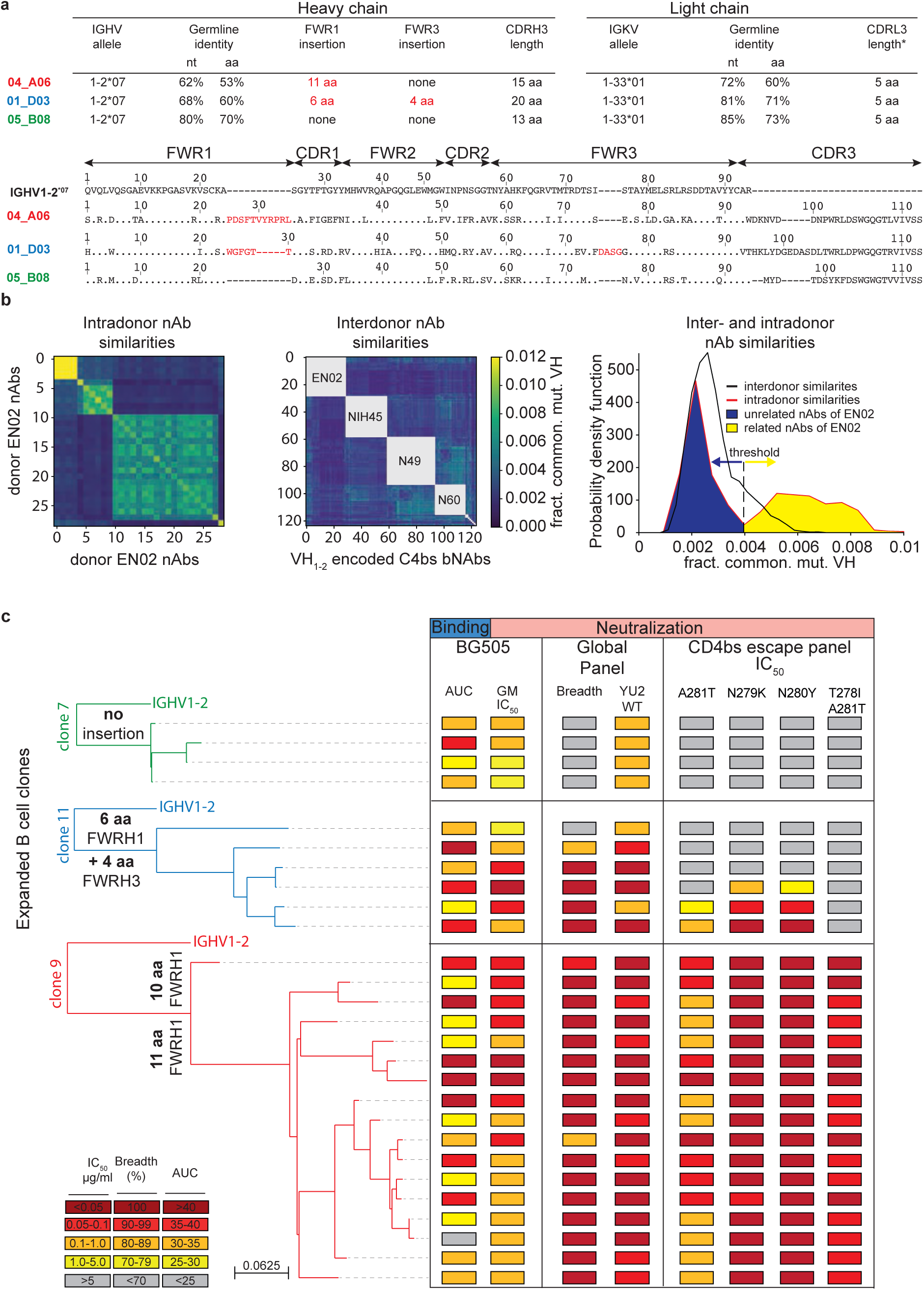
bNAb coevolution with overlapping specificity for the CD4bs. Phylogenetic and functional analyses reveal co-existence of genetically-distinct CD4bs bNAbs in donor EN02. **a**, Antibody characteristics and alignment of 04_A06, 01_D03, and 05_B08 heavy chains to germline IGHV1-2 gene. **b**, Heatmaps illustrating the V_H_ sequence similarities of isolated antibodies within (intradonor) and between (interdonor) donors in shades from blue to yellow. Donors of other VH1-2-encoded CD4bs bNAbs used as comparators are displayed in the center of each grey square. Histogram outlining intra- and interdonor similarity distributions. Red line indicates intradonor similarity distribution for donor EN02; black line indications interdonor similarity distribution between donors EN02 and VH1-2-encoded CD4bs bNAbs from other donors; dashed line indicates the threshold for identification of clonally-related sequences. Shaded areas under the intradonor similarity distribution graph indicate non-clonal (blue) and clonal (yellow) sequences. **c**, Maximum-likelihood phylogenetic trees of isolated bNAbs encoded by IGHV1-2 (green, blue, red). Antibodies were tested for binding (mean AUC) to BG505_SOSIP.664_ and neutralizing activity (GeoMean IC_50_ and breadth) against a HIV-1 global pseudovirus panel^64^ and common CD4bs escape variants. Samples were tested in duplicates. Reference antibodies were validated for functionality in neutralization assays against the global HIV-1 pseudovirus panel, and only those with IC₅₀/IC₈₀ values deviating less than threefold from CATNAP reference data^128^ were included.

Members of the identified expanded B cell clones encoding VH1-2-derived nAbs were tested for BG505_SOSIP.664_ binding and neutralizing activity against the full HIV-1 global pseudovirus panel^64^ (Fig. 2c). Competed binding activity by other IGHV1-2-encoded CD4bs bNAbs determined for representative B cell clone members suggested overlapping CD4bs epitope specificity (Extended Data Fig. 3c). Except for one mAb, all evaluated antibodies bound soluble BG505_SOSIP.664_ and all mAbs neutralized the global pseudovirus panel. Potency and breadth were associated with mutation frequency and heavy chain amino acid insertions (Fig. 2c, Extended Data Fig. 3d).

Members of clones 9 and 11 achieved up to 100% breadth, while clone 7 members reached only 50% against the global pseudovirus panel (Fig. 2c, Supplementary Table 3). These results were confirmed for representative members of each clone when tested against ≥245 strains (Extended Data Fig. 3d). Testing against VRC01-class escape variants of the HIV-1 strain YU2 showed high activity only in clone 9 members with 10 or 11 amino acid insertions in the FWRH1, while members of the remaining clones neutralized few or no variants (Fig. 2c). To determine whether the 11-amino acid insertion mediates restriction of viral escape, we transferred the insertion from 04_A06 into representative members of the other isolated clones and other IGHV1-2-encoded CD4bs bNAbs (Extended Data Fig. 3e-g). Incorporating this insertion into 05_B08 and VRC07 (VRC07_FWR-Ins_) maintained activity against the global HIV-1 and/or 119 multiclade panel and enabled these chimeric antibodies to overcome VRC01-class escape variants *in vitro* (Extended Data Fig. 3e,f). However, VRC07_FWR-Ins_ failed to demonstrate antiviral activity against rebound viruses emerging during VRC07 monotherapy in HIV-1_YU2_-infected mice *in vivo* (Extended Data Fig. 3g). Additionally, unlike a representative member of clone 9 (04_A06) from which the 11-amino acid insertion was transferred, VRC07_FWR-Ins_ displayed high autoreactivity (Extended Data Fig. 4a). These findings suggest that although the identified B cell clones share overlapping CD4bs specificities, they differ in unique genetic adaptations, such as the development of ultralong FWRH1 amino acid insertions, which contribute to the restriction of VRC01-class viral escape.

### A highly active CD4bs bNAb with remarkable breadth and potency

From clone 9, we selected antibody 04_A06 for further characterization. First, we determined the binding profiles of 04_A06 and VRC01-class CD4bs bNAbs against Env-derived proteins. Whereas both 04_A06 and VRC01-class antibodies were reactive to YU2_gp120_, 93THO57_gp120/kif_, and YU2_gp140_, 04_A06 did not bind to the V1–V3 loop-deficient resurfaced stabilized gp120 core 3 (RSC3) that was applied to selectively enrich VRC01-class antibodies in bait protein-based single cell sorting strategies^37,53^ (Extended Data Fig. 4b). These results implied a distinct mode of epitope recognition for 04_A06 compared to other VRC01-class bNAbs.

Next, we determined the antiviral activity of 04_A06 against diverse reference HIV-1 pseudovirus panels comprising a total of 337 viral strains and against 50 outgrowth culture-derived primary HIV-1 isolates (Fig. 3, Extended Data Fig. 4, Supplementary Table 4-8). 04_A06 neutralized 100% of the 12-strain HIV-1 global panel^64^ with higher potency (GeoMean IC_50_/IC_80_ = 0.038/0.184 µg/ml) and/or breadth than clinically-advanced bNAbs and previously identified CD4bs bNAbs (Fig. 3a, Extended Data Fig. 4c, Supplementary Table 4). We extended the evaluation of 04_A06 against well-established large multiclade pseudovirus panels that cover difficult-to-neutralize (tier-2 and tier-3) viruses, major genetic HIV-1 subtypes, and circulating recombinant forms.

**Figure 3:**
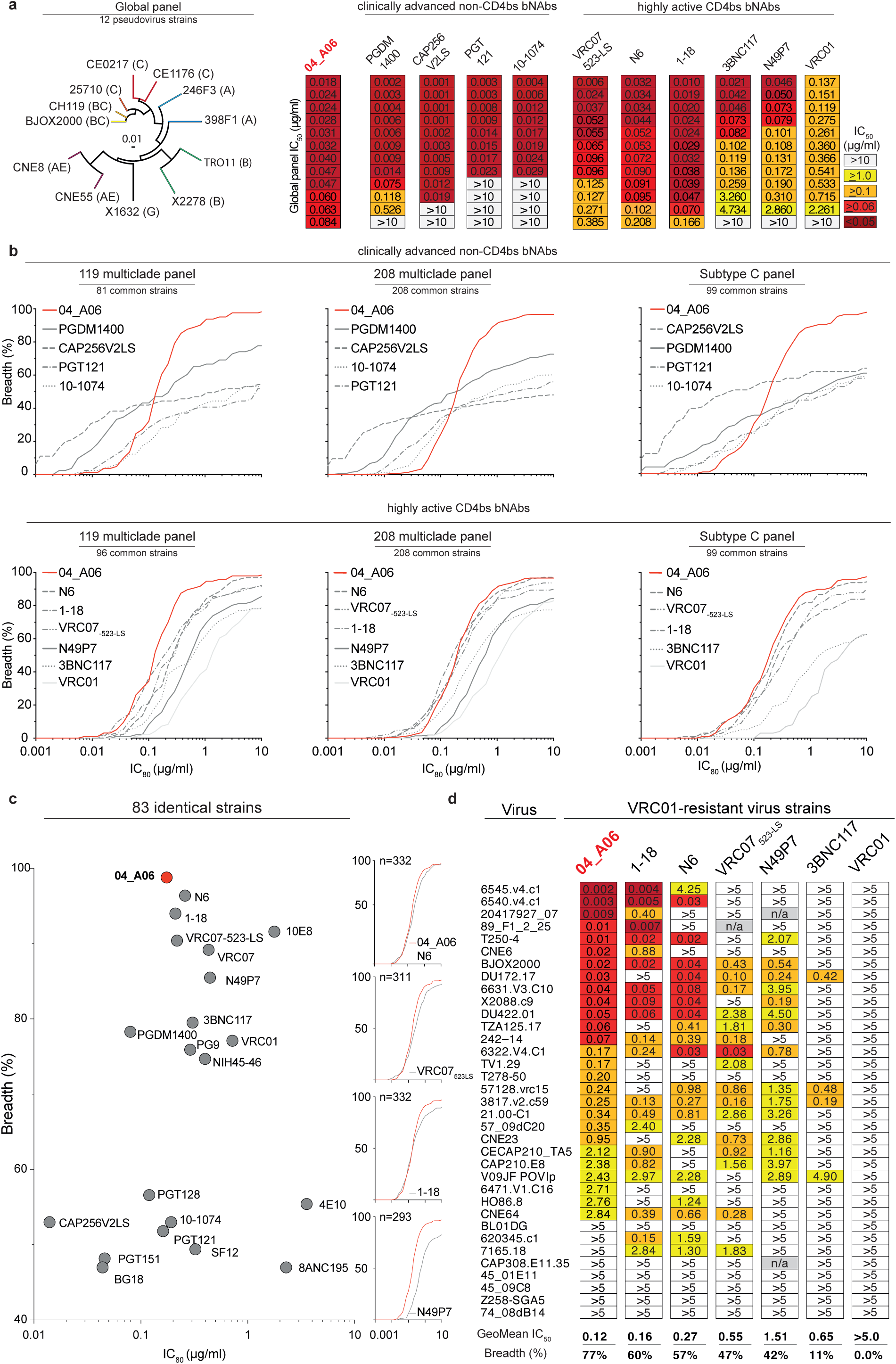
04_A06 demonstrates remarkable neutralizing potency and breadth. **a**, Illustration of the phylogenetic distribution of Env-pseudotyped viruses of the 12-strain global panel^64^. Neutralizing profile of 04_A06 against the global panel compared to non-CD4bs bNAbs currently investigated in advanced clinical trials and to highly-active CD4bs bNAbs. Illustrated IC_50_ values for each bNAb are sorted by increasing value. **b**, Neutralization coverage (%) and potency (IC_80_) of 04_A06 against multiclade panels of 119^74^ and 208^75^ pseudoviruses, and a Clade C pseudovirus panel. Data are shown for identical virus strains from each panel for which reference neutralization data was available. **c,** Dot plot illustrating the neutralizing activity (IC_80_, breadth) of 04_A06 compared with anti-HIV-1 bNAbs against 83 pseudoviruses. Neutralization data of bNAbs N6, 1-18, and 04_A06 were determined in the same laboratory. Curve graphs compare neutralizing activity of 04_A06 to current highly active CD4bs bNAbs (N6, VRC07_523-LS_, 1-18, N49P7) against panels of 332, 311, 332, or 293 pseudoviruses. **d**, Neutralization profile (IC_50_, breadth) of 04_A06 compared with highly active CD4bs bNAbs against a panel of 37 pseudoviruses resistant to CD4bs bNAbs. Data for reference bNAbs obtained from the CATNAP database^128^ if available. Otherwise bNAbs were tested in parallel to 04_A06. Reference antibodies were validated for functionality in neutralization assays against the global HIV-1 pseudovirus panel, and only those with IC₅₀/IC₈₀ values deviating less than threefold from CATNAP reference data^128^ were included. Breadth (%) was defined applying a cut-off of ≤10 µg/ml (a-c) and ≤5 µg/ml (d). Samples were tested in duplicates in single experiments. Geometric mean neutralization data are presented.

Notably, against these virus panels 04_A06 exhibited comparable neutralizing activity to previously-described CD4bs bNAbs (Fig. 3b,c, Supplementary Table 5-7). 04_A06 displayed high potency (GeoMean IC_50_/IC_80_ = 0.037/0.135 µg/ml) and breadth (98.3%/98.3%) against a 119-strain multiclade panel^51,54,57,74^ (Fig. 3b, Supplementary Table 5). In addition, 04_A06 demonstrated remarkable potency and neutralization breadth (GeoMean IC_50_/IC_80_=0.077/0.198µg/ml; Breadth=98.6%/96.6%) when tested against a 208-strain multiclade panel^75^ (Fig. 3b, Supplementary Table 6). Furthermore, 04_A06 exhibited high activity against a 100-strain clade C panel^76^ (GeoMean IC_50_/IC_80_=0.057/0.192 µg/ml; Breadth=99%/98%) (Fig. 3b, Supplementary Table 7). Across all 337 HIV-1 pseudovirus reference strains evaluated, only 5 or 9 strains were resistant against 04_A06 at an IC_50_/IC_80_ ≥10 µg/ml (GeoMean IC_50_/IC_80_=0.059/0.176 µg/ml, Breadth=98.5/97.3%) (Fig. 3b,c, Supplementary Table 5-7). Pseudoviruses derived from plasma single genome sequencing (SGS) *env* sequences of donor EN02 that comprised rare amino acid residues or insertions, exhibited resistance to 04_A06 as well as to other CD4bs-targeting bNAbs (Extended data Fig. 4e,f). We also determined the activity of 04_A06 against replication-competent donor-derived outgrowth viruses, which are more challenging to neutralize than pseudoviruses^77^. Against viruses obtained from 50 PLWH, 04_A06 demonstrated levels of breadth and/or potency (GeoMean IC_50_/ IC_80_=0.45/1.736 µg/ml, Breadth=94%/88%) comparable to highly-active CD4bs bNAbs. (Extended Data Fig. 4d, Supplementary Table 8). Finally, we investigated the activity of 04_A06 against a panel of 35 strains with pronounced resistance to VRC01-class antibodies (e.g., full resistance to VRC01)^33–35,37,78^ (Fig. 3d). Notably, whereas the near-pan-neutralizing CD4bs bNAbs 1-18^36^ and N6^34^ neutralized only 57% and 60% of viruses, respectively, 04_A06 achieved a breadth of 77% with higher potency (GeoMean IC_50_=0.12 µg/ml).

Through testing against large panels of pseudo- and replication-competent viruses, we demonstrated that 04_A06 is a remarkable CD4bs bNAb with substantial activity against even highly resistant viral strains.

### Ultra-long 11-amino acid insertion facilitates the quaternary binding mode of 04_A06 to the CD4bs and adjacent gp120 protomer

To elucidate mechanisms of Env recognition, we determined single particle cryo-EM structures of a BG505_SOSIP.664_ trimer^67^ both unbound (2.9 Å) and in complex with Fabs from representative bNAbs from each expanded B cell clone: 04_A06 (3.8 Å), 01_D03 (3.2 Å), or 05_B08 (3.3 Å) (Fig. 4a, Extended Data Fig. 5, Extended Data Fig. 6a). The 04_A06 complex also included a Fab from PGDM1400, a broader and more potent variant of the V1V2 bNAb PGT145^79^. In processing this dataset, particles were separated into classes with and without PGDM1400, thereby also resulting in a Fab-Env complex structure of this clinically-relevant antibody (Extended Data Fig. 5).

**Figure 4:**
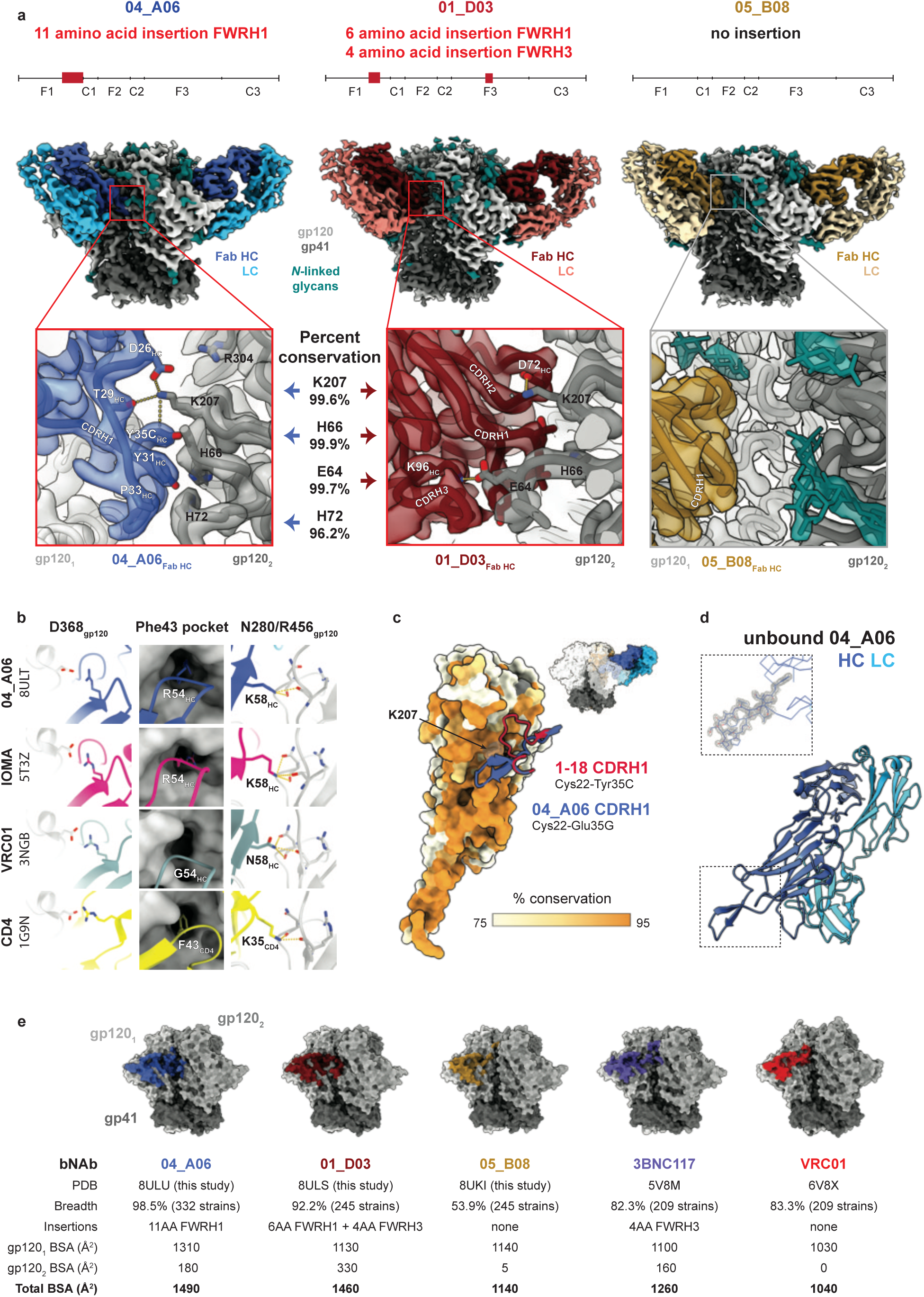
Ultra-long 11-amino acid insertion facilitates quaternary binding mode of 04_A06 a, EM maps showing side views of 04_A06 (blue), 01_D03 (red) and 05_B08 (gold) bNAb Fabs in complex with BG505_SOSIP.664_ Env trimers. Insets illustrate a close up of bNAb interactions with the adjacent gp120 protomer (gp120_2_). **b,** Canonical interactions of CD4 and CD4bs bNAbs with Env: D368_gp120_, Phe43 pocket, and N280_gp120_ / R456_gp120_. See Extended Data Fig. 6b for interactions with additional bNAbs including 01_D03 and 05_B08. **c,** Interactions between the CDRH1s of 04_A06 and 1-18, shown as blue and red cartoon representations, with the secondary (gp120_2_) Env protomer, shown as a surface colored by percent conservation. **d,** Crystal structure of unbound 04_A06 Fab. Inset highlights the 04_A06 CDRH1, with electron density contoured at 1.5!. **e,** Surface area buried by VRC01-class bNAbs on primary (gp120_1_) or secondary (gp120_2_) Env protomers.

All three CD4bs bNAbs exhibited canonical gp120 contacts made by CD4 and other CD4-mimetic bNAbs^39,40,80^ (Fig. 4b, Extended Data Fig. 6b). In each structure, R71_HC_ contacts D368_gp120_, a feature typical of VRC01-class antibodies and a hallmark of CD4-mimetic bNAbs^40^. Additionally, the V_H_1-2-encoded S54_HC_ is mutated to R54_HC_ in 04_A06 and Y54_HC_ in 01_D03, both of which insert into a hydrophobic cavity on gp120 analogous to the interactions of CD4 residue F43_CD4_^81^. These interactions are similar to the interactions made by R54_HC_ of the VH1-2 encoded, non-VRC01-class bNAb, IOMA^80^ and Y54_HC_ of the VRC01-class bNAb N6^34^ (Fig. 4b, Extended Data Fig. 6b). Additionally, the V_H_1-2–encoded N58_HC_ is mutated to K58_HC_ in both 04_A06 and IOMA, mimicking the interaction of K35_CD4_ with Env residues N280_gp120_ and R456_gp120_ (Fig. 4b). Although 04_A06 features a five-amino acid-long CDRL3 characteristic of VRC01-class bNAbs, it also shares key somatic hypermutations and structural determinants with IOMA shaping recognition of the CD4bs. Thus, 04_A06 possesses features representative of both VRC01-class and IOMA-like CD4bs bNAbs.

The presence and length of insertions in the three representative bNAbs correlated with the formation of interprotomer contacts (Fig. 2a, 4a). 04_A06 and 01_D03, but not 05_B08, contacted the adjacent gp120 protomer, a feature seen in other potent bNAbs such as 1-18^36^ and 3BNC117^82^ (Fig. 4a). Contacts between 04_A06 and the adjacent protomer were mediated exclusively via a protruding CDRH1 in 04_A06, a consequence of the 11-residue FWRH1 insertion (Fig. 4a), which also contacted the primary protomer (Extended Data Fig. 6c). On the adjacent protomer, 04_A06 interacted with K207_gp120_ (99.6% conserved) (www.hiv.lanl.gov and^83^) and formed a potential electrostatic interaction with D26_HC_. D26_HC_ is also positioned adjacent to R304_gp120_ (93.5% conserved), while residues within the tip of the 04_A06 CDRH1 (Y31_HC_-Y35C_HC_) are in close proximity to residues H66_gp120_ and H72_gp120_ (99.9%, 96.2% conserved, respectively) (Fig. 4a). Together, these residues could form interactions that permit tolerance to structural variability within the CD4bs on the primary Env protomer (e.g., addition of a potential N-linked glycosylation site at N279_gp120_) (Fig. 2c). The extended CDRH1 of the CD4bs bNAb 1-18 also contacts K207_gp120_ on the adjacent protomer but mainly interacts with less conserved V3 residues^36^, whereas the CDRH1 of 04_A06 extends toward a more conserved gp120 region, likely contributing further to 04_A06’s enhanced neutralization profile (Fig. 3d,4c). In contrast to 04_A06’s CDRH1-mediated interprotomer contacts, mAb 01_D03, with its 6-residue FWRH1 insertion, 4-residue FWRH3 insertion, and 20-residue CDRH3, contacts the adjacent protomer with each of its heavy chain CDRs and makes potential electrostatic interactions with K207_gp120_ and E64_gp120_ (99.6% and 99.7% conserved) (Fig. 4a). In the 01_D03-complexed Env structure, but not in the unbound

BG505 SOSIP structure, EM density was observed for residues 57-65_gp120_ on the adjacent protomer, suggesting that this normally disordered loop becomes stabilized upon antibody binding (Extended Data Fig. 6d).

To determine whether the conformation of the extended CDRH1 of 04_A06 is pre-organized for binding, we solved a 1.75Å crystal structure of unbound 04_A06 Fab (Fig. 4d, Supplementary Table 9). The structure, which does not appear to be influenced by crystal contacts (Extended Data Fig. 6e), revealed a well-ordered antibody combining site and CDRH1 that resembled the conformation of the combining site in the Env-bound Fab (root mean square deviation, rmsd = 0.64 Å; 235 C⍺ atoms), consistent with a lock-and-key mechanism of binding^84^ (Fig. 4d, Extended Data Fig. 6f). This is in contrast to the induced fit mechanism of binding by the extended CDRH1 of VRC-CH31, which is unresolved in crystal structures in complex with gp120^85^, but ordered in a VRC-CH31–SOSIP Env structure in which the CDRH1 is stabilized by interactions with the adjacent protomer^86^. The pre-organized antibody combining site of 04_A06 likely leads to a more favorable interaction with Env due to a lower entropic cost of binding^87^.

In addition to contacting highly conserved residues, the insertions and interprotomer contacts of 04_A06 and 01_D03 contribute to an increased amount of surface area buried on Env by antibody binding. These insertions enable 04_A06 and 01_D03 to bury more surface area on Env than 05_B08 or other VRC01-class bNAbs (Fig. 4e), presenting another possible mechanism contributing to breadth, potency, and resistance to escape^36,88^. Consistent with this and in common with 04_A06, 01_D03 exhibits greater potency and breadth than 3BNC117 and VRC01 (GeoMean IC_50_/IC_80_=0.052/0.187 µg/ml, Breadth=92%/88%; 245 strains) (Fig. 4e, Extended data Figure 4d).

Our 04_A06 complex structure also included PGDM1400, a more broad and potent somatic variant of PGT145^79^ (Extended Data Fig. 7a). The PGDM1400 Fab binds asymmetrically to the trimer apex with a stoichiometry of one Fab per trimer (Extended Data Fig. 7b). Its 34-residue CDRH3 is inserted down the trimer symmetry axis and contacts protein residues and the N160_gp120_ N-glycan of all three protomers, resulting in well-resolved glycan density for the core pentasaccharide of each N160_gp120_ glycan and additional glycans in some cases (Extended Data Fig. 7c). Mass spectrometry analysis indicated that the N160_gp120_ glycan is under-processed, containing a mixture of high-mannose, hybrid, and complex-type N-glycans^89^. In our structure, density was not observed for a core fucose, a component of complex-type N-glycans^80^, nor could clear density be discerned much beyond the core pentasaccharide, the latter a likely consequence of limited resolution, glycan compositional heterogeneity, and/or glycan flexibility. However, a surface electrostatic calculation revealed electropositive patches that could accommodate, or interact with, negatively-charged sialic acid residues on complex-type glycans^90,91^ (Extended Data Fig. 7c).

This observation is particularly notable for surfaces on PGDM1400 that interact with the N160_glycan2_ and N160_glycan3_ relative to analogous interactions with PGT145^82^ (Extended Data Fig. 7c,d). Additionally, PGDM1400 appears to stabilize the core pentasaccharide of the N160_glycan3_, whereas this glycan was proposed to be inhibitory to the binding of PGT145^82^. The ability to accommodate and/or interact with the three N160_gp120_ glycans on an Env trimer, whether or not processed beyond high-mannose carbohydrates, likely contributes to the enhanced breadth and potency of PGDM1400. Our structure recapitulates key molecular interactions between PGDM1400 and Env recently reported^92^. For example, K169_gp120_ interacts with Tys100F_HC_, an Asp-Asp-Asp motif at the tip of the PGDM1400 CDRH3 forms electrostatic interactions with R166_gp120_ of all three protomers, and extensive glycan density was observed at N160_gp120_ on all protomers.

### 04_A06#restricts viral escape and fully suppresses viremia in humanized mice

Determining the potential for viral escape from nAbs provides critical insights for development of antibody-based prevention and treatment strategies. First, we assessed the antiviral activity of 04_A06 against a library of known HIV-1 Env escape mutations in the HIV-1 BG505_T332N_ background^1,2,7,8,47,50,51,60,79,93–95^. Notably, none of the known viral escape mutations tested, including those located in the gp120 loop D that typically interfere with CD4bs bNAbs (N279K and A281T)^2,36,50,96,97^, conferred resistance against 04_A06 (Extended Data Fig. 8a). Next, we applied a lentivirus deep mutational scanning (DMS) platform^98–100^ to comprehensively measure how all functionally tolerated Env_BF520_ mutations affect neutralization by 04_A06 and other CD4bs reference bNAbs (Extended Data Fig. 8b). Env mutations in loop D caused escape from bNAbs N6LS (A281W and A281T), VRC07-523-LS (N279K), and 3BNC117 (N279R/E/K and A281W/F/D/I/V). Additionally, mutations in the beta23/V5 loop caused escape from N6LS (G451D) and 3BNC117 (R456W/S and G471). Amino acid substitutions potentially reducing neutralization sensitivity of antibody 1-18 were identified at sites 198_gp120_, 203_gp120_, and 206_gp120_ near the N197_gp120_ glycosylation motif and V2 loop, at sites 428-430_gp120_ in the beta20/beta21 regions, and at sites 47_gp120_, 474_gp120_, and 476_gp120_ in the beta23/V5 loop regions (Extended Data Fig. 8b). Interestingly, DMS did not identify any single mutations to Env_BF520_ that strongly escaped 04_A06, with just a few mutations (e.g., V164W and Q428Y) possibly slightly reducing neutralization sensitivity. The lack of prominent escape in DMS sets 04_A06 apart from other highly broad and potent CD4bs bNAbs (Extended Data Fig. 8b).

To investigate the *in vivo* antiviral activity of 04_A06, we infected humanized mice with HIV-1_YU2_ and subcutaneously administered 1 mg of 04_A06 followed by 0.5 mg of antibody twice a week for up to 12 weeks (Fig. 5). PBS-treated mice utilized as untreated controls showed maintained viremia for ≥12 weeks (Extended Data Fig. 3g). Administration of the CD4bs bNAbs VRC01 and VRC07 (the parental form of VRC07_523-LS_) resulted only in a transient reduction of viremia (0.54/0.8 log_10_ copies/ml), generally followed by viral rebound within 3 weeks of treatment initiation (Fig. 5a). Similarly, monotherapy with N49P7 and N6 led to temporary viral load reductions in the majority of animals (6/7 for N49P7; 3/6 for N6), with rebound occurring between 14- and 28-days post-treatment initiation (Extended Data Fig. 10a). In contrast and despite similar sensitivity of the HIV-1_YU2_ challenge stock to all administered bNAbs, 04_A06 therapy decreased the mean HIV-1 viral load by 1.96 log_10_ copies/ml and resulted in maintained suppression of viremia to levels below the lower limit of quantification (LLQ <784 or 451 copies/ml) in all animals (19/19 mice) without the occurrence of viral rebound (Fig. 5a, Extended Data Fig. 10a, Supplementary Table 10). To investigate the development of viral escape mutations, we performed envelope single genome sequencing from plasma^101^ obtained 4 weeks after treatment initiation. Viral rebound in VRC01-or VRC07-treated mice was associated with the accumulation of mutations in the Env loop D and/or beta23/V5 loop^36^ (Fig. 5b,e, Supplementary Table 10). These mutations mainly affected Env residues N279_gp120_, N280_gp120_, A281_gp120_ and G459_gp120_, thereby decreasing sensitivity to VRC01 and VRC07 but not to 04_A06. Moreover, N49P7 and N6 monotherapy selected for mutations within the loop D at Env residues N276_gp120_, A281_gp120_ and K282_gp120_ (Extended Data Fig. 10b, Supplementary Table 10). A281_gp120_ represents a key contact residue for N49P7^33^ and DMS in this study as well as previous reports^36^ identified the A281T_gp120_ substitution to mediate resistance against N6. In contrast, in the 04_A06-treated mice not yet fully suppressed at week 4, we found no selection of mutations affecting 04_A06 sensitivity (Fig. 5b,e, Supplementary Table 10). Finally, to determine 04_A06 activity against VRC01-resistant viral escape variants *in vivo*, we administered 04_A06 to VRC01-pretreated animals following rebound. To maintain VRC01 selection pressure, we continued VRC01 therapy after initiation of 04_A06 therapy (1 mg loading dose followed by 0.5 mg twice weekly for each antibody). Mice receiving a double dose of VRC01 served as control group (1 mg of VRC01 as loading dose followed by 1 mg twice weekly). While the increased dose of VRC01 in the control group had no effect on viral loads, addition of 04_A06 to the treatment regimen led to complete suppression of viremia for 8 weeks in all VRC01 pre-treated animals (Fig. 5c).

**Figure 5:**
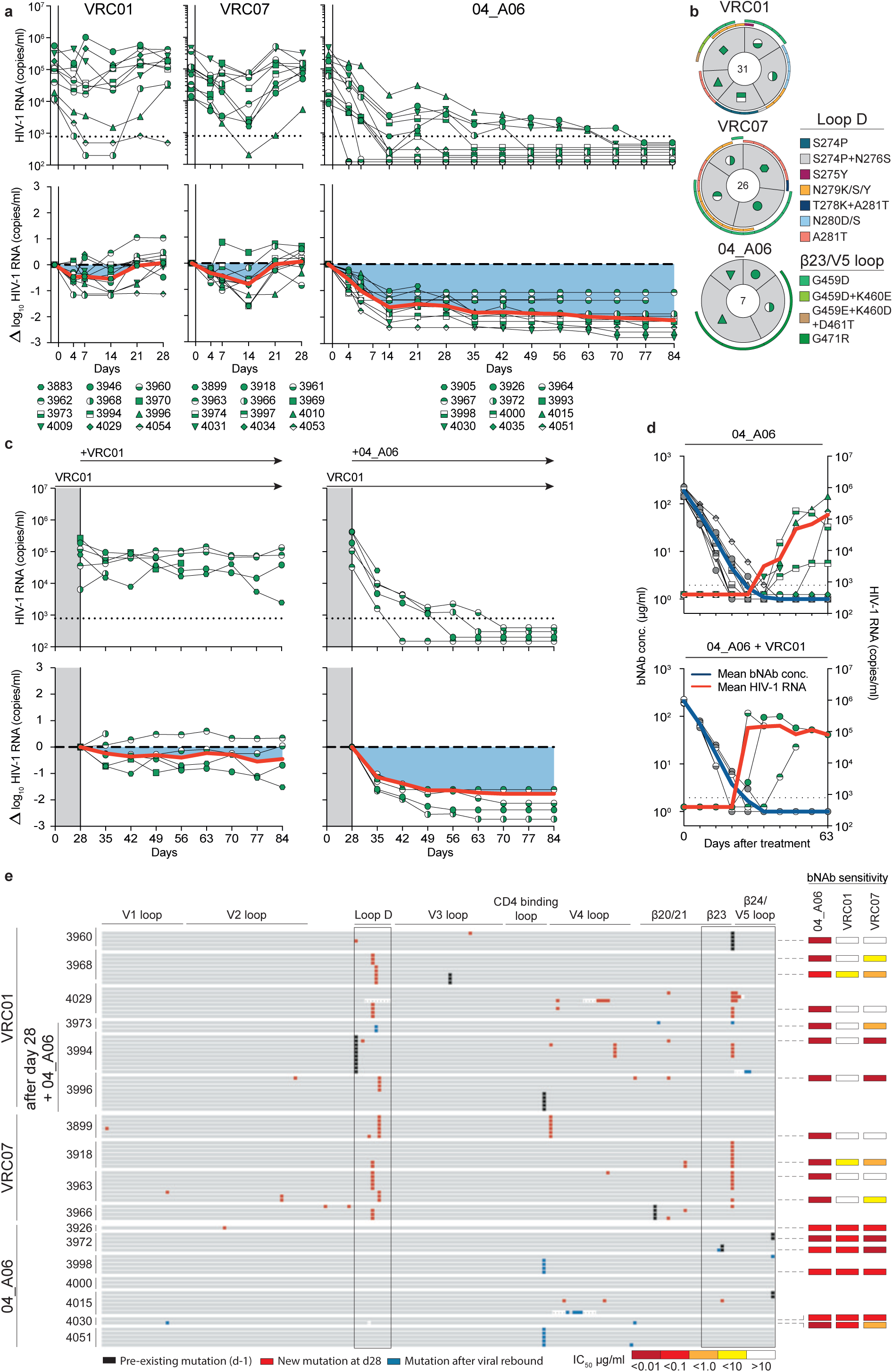
Maintained suppression of viremia and restriction of VRC01-class viral escape *in vivo*. **a**, Investigation of the antiviral activity of 04_A06, VRC01, and VRC07 monotherapy in HIV-1_YU2_-infected humanized mice (NRG). Graphs illustrate the absolute HIV-1 RNA plasma copies/ml (top) and relative log_10_ changes from baseline viral loads (bottom) after initiation of bNAb therapy. Dashed lines (top graphs) indicate the lower limit of quantitation of the qPCR assay (LLQ) (784 copies/ml). Red lines display the average log_10_ changes compared to baseline viral loads (day −1). **b**, Analyses of single HIV-1 plasma *env* sequences from HIV-1_YU2_-infected humanized mice obtained on day 28 after bNAb treatment initiation. Total number of analyzed sequences is indicated in the center of each pie chart. Mice are labeled according to icon legends in **a**. Colored bars on the outside of the pie charts indicate mutations in Loop D and beta23/V5 loop. **c**, Sequential treatment with 04_A06 in HIV-1_YU2_-infected humanized mice following viral rebound during VRC01 monotherapy (from panel **a**). This approach included maintaining VRC01 monotherapy while integrating 04_A06 in the treatment regimen. Mice treated with a double dose of VRC01 (1mg) were included as a control group. Dashed lines in (top graphs) indicate the qPCR assay LLQ (784 copies/ml). Red lines display average log_10_ changes compared to baseline viral loads (day 28). **d**, HIV-1 viral loads (right y-axis) and plasma bNAb concentrations (left y-axis) after treatment interruption on day 84. Blue line indicates the mean plasma bNAb concentration; red line displays geomean HIV-1 viral loads. Dashed lines indicate the qPCR assay LLQ (784 copies/ml). Mice are labeled according to icon legends in **a**. Grey icons refer to measured antibody plasma levels; green icons indicate viral loads. **e**, Alignment of plasma SGS-derived *env* sequences identified from day −1 (black bars), day 28 (red bars), and after viral rebound (blue bars; panel **d**) and sensitivity of the respective pseudovirus.

To determine whether waning antibody levels after treatment interruption could promote 04_A06 escape, we discontinued therapy and monitored mice for 9 weeks (Fig. 5d,e). The median duration to viral rebound was 38.5 days and rebound occurred after plasma antibody levels decreased below 3 µg/ml (04_A06 + VRC01) or were undetectable (04_A06) (LLQ < 1 µg/ml). Of 13 mice, viremia did not rebound in 3 mice until their demise or the end of the 9-week interval, potentially due to graft failure. Single genome sequencing of plasma rebound virus *env* genes, together with limited functional characterization of a subset of derived pseudoviruses, did not reveal consistent selection of specific amino acid mutations conferring resistance to 04_A06 across all animals (Fig. 5e, Supplementary Table 10 - 11). We conclude that 04_A06 monotherapy effectively mediates long-term control of viremia in HIV-1_YU2_-infected humanized mice and can restrict and overcome VRC01-class viral escape *in vitro* and *in vivo*.

### *In silico* analyses predict potential of 04_A06 for HIV-1 prevention

To assess the potential of 04_A06 for prevention strategies, we determined its activity against a representative panel of AMP trial-derived HIV-1 pseudoviruses (Fig. 6a, Extended Data Fig. 9, Supplementary Table 12), allowing assessment of antiviral activity against contemporaneous-circulating HIV-1 strains. These pseudoviruses were based on *env* sequences observed after HIV-1 infection and represent infection in the absence of selection pressure (placebo arms, HVTN 703/HPTN 081 and HVTN 704/HPTN 085) or breakthrough infections in the presence of VRC01 (VRC01 arm of HVTN703/HPTN 081 trial). Most pseudoviruses belonged to clade C (HVTN 703/HPTN 081) or blade B (HVTN704/HPTN 085). Antibody 04_A06 displayed high levels of potency and breadth against viruses derived from both the placebo group (GeoMean IC_50_/IC_80_=0.07/0.26 µg/ml; Breadth=100%/98%, at ≤10 µg/ml) and the HVTN703 treatment group (GeoMean IC_50_/IC_80_=0.10/0.31 µg/ml; Breadth=97%/94%, at ≤10 µg/ml) (Fig. 6a, Extended Data Fig. 9a,b). In the AMP trials, a viral sensitivity threshold of IC_80_<1 µg/ml was associated with VRC01-mediated protection against HIV-1 infection^9,102^. When applying this threshold, 04_A06 exhibits high activity against viruses generated from the placebo-(GeoMean IC_80_=0.20 µg/ml, at <1 µg/ml) and the HVTN703 treatment groups (GeoMean IC_80_=0.18 µg/ml at <1 µg/ml) (Fig. 6a, Extended Data Fig. 9a,b). For example, whereas VRC01 neutralized only 33% and 9.3% of the AMP placebo and HVTN703/HPTN 081 treatment group pseudoviruses at this threshold, 04_A06 achieved 87% or 74% breadth. Notably, 04_A06 also remained active against pseudoviruses generated from breakthrough infections of the HVTN703 treatment group that exhibited high resistance to VRC01 (Extended Data Fig. 9b, Supplementary Table 12).

**Figure 6:**
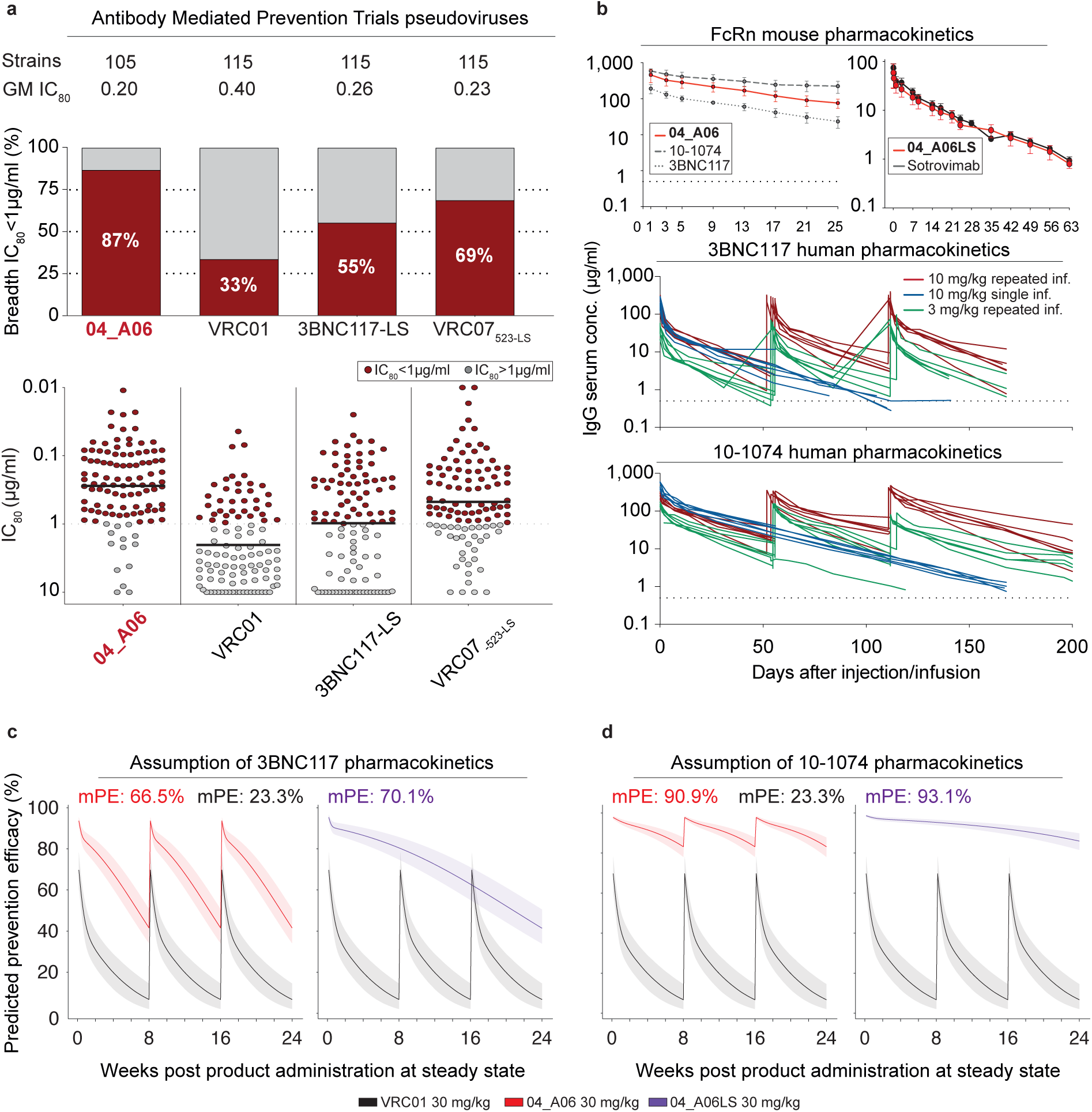
Enhanced antiviral activity against AMP trial viruses and high HIV-1 prevention efficacy. **a**, Illustration of the neutralizing activity of 04_A06 compared to CD4bs bNAbs against representative pseudoviruses generated from the placebo arms of the two AMP trials: HVTN703/HPTN 081 in predominantly Clade B regions and HVTN704/HPTN 085 in predominantly Clade C regions^9,102,112^. Bar graphs display the breadth (%) at the established threshold of protection (IC_80_ <1 μg/ml). Dot plots indicate the IC_80_s of bNAbs against the individual pseudoviruses. Virus strains neutralized at the established threshold of protection (IC_80_ <1 μg/ml) are depicted in red, whereas strains not neutralized at this threshold are shown in grey. Data for reference bNAbs were retrieved from the CATNAP database^128^ and^112^. Antibodies were tested in duplicates. The center line in dot plots depicts the geometric mean. **b**, Serum concentrations of 3BNC117 and 10-1074 from a human clinical trial^106^ and *in vivo* pharmacokinetic profile of 04_A06 and 04_A06LS measured in hFcRn transgenic mice. For determination of the PK profile, mice received either a fixed dose of 0.5 mg of 04_A06 or 5 mg/kg of 04_A06LS. Serum concentrations (μg/ml) of 04_A06 (top), 3BNC117 (middle), and 10-1074 (bottom). Participants of the human clinical trial received either a single infusion of 3BNC117 and 10-1074 (10 mg/kg of each bNAb, blue), or three infusions of each bNAb (3 mg/kg, green; 10 mg/kg, red). Mice received a single intravenous injection of 0.5 mg of 04_A06. Serum concentrations were determined in duplicates by ELISA. Data of mice experiments are represented as mean ± SD. **c**, Predicted HIV-1 prevention efficacy (PE) of 04_A06 or VRC01 over time against AMP viruses. Left: PE of 04_A06 (red) 04_A06LS (purple) assuming of 3BNC117 PKs^106^. Right: PE of 04_A06 and 04_A06LS assuming 10-1074 PKs. PE was calculated over time after three 8-weekly infusions of 04_A06 or a single infusion of 04_A06LS (all at 30 mg/kg). For LS versions of antibodies, predictions were made assuming that 04_A06-LS has a 2.5-fold longer half-life than 04_A06. Black curves indicate the VRC01 PE in the AMP trials. Solid lines display the median and shaded areas the 95% prediction interval.

Next, we modeled the HIV-1 prevention efficacy (PE) of 04_A06 by applying the predicted serum neutralization 80% inhibitory dilution titer (PT_80_) recently established as a correlate of protection^102^ (Extended Data Fig. 9c). This biomarker integrates the *in vitro* neutralization potency (IC_80_) and pharmacokinetic (PK) profile of a bNAb, thereby quantifying neutralizing activities against exposure viruses at variable time points. Since PK data are required for *in silico* PE estimation, we first investigated the PK profile of 04_A06 in transgenic mice that express the human neonatal Fc receptor (hFcRn), which closely mirror antibody PK behavior in humans^103–105^ (Fig. 6b). Following a single intravenous injection, 04_A06 displayed a PK profile with a maximum concentration and rate of decline between those observed for bNAbs 3BNC117 and 10-1074^106^. The LS-engineered variant of 04_A06 closely resembled the PK profile of Sotrovimab^107^, an anti-SARS-CoV-2 mAb with enhanced hFcRn binding and extended systemic half-life life. We hypothesized that the PK behavior of 04_A06 parallels those of 3BNC117 and 10-1074 observed in humans and modeled the PT_80_ and PE of 30 mg/kg 04_A06 under the assumption of either 3BNC117-like or 10-1074-like PKs^106^ (Fig. 6b-d). In addition, we incorporated PK predictions for the LS form of 04_A06 (04_A06LS) considering a conservative 2.5-fold increase in the elimination half-life^43^. Simulations were performed based on PK modeling for a period of 24 weeks with bNAb infusions in 8-week intervals for non-LS antibodies^102,108^. The time-averaged mean predicted prevention efficacy (mPE) of 30 mg/kg 04_A06 assuming 3BNC117 or 10-1074 PKs was 66.5% or 90.9%, respectively, exceeding the 23.3% mPE of VRC01 given every 8 weeks (Fig. 6c). Simulation of predictions for a single administration of 04_A06_LS reached 70.1% or 93.1% mPE over a period of 6 months. In conclusion, 04_A06 demonstrates remarkable antiviral efficacy against pseudoviruses derived from the AMP trials, establishing this bNAb as a promising antibody candidate for future prevention strategies.

## Discussion

Clinical trials have highlighted the potential of anti-HIV-1 bNAbs as effective and generally well-tolerated options for prevention and treatment of HIV-1 infection^1,2,5–7,9,47,95^. However, the identification of improved bNAbs and delineation of their developmental pathways remains essential to fully exploit bNAb immunotherapy opportunities and advance the development of an effective HIV-1 vaccine^102^. A major challenge in this endeavor is the extraordinary rarity of bNAbs, which are found in only a small fraction of PLWH mounting a broad serum neutralization response^58,59,109,110^. Although various anti-HIV-1 bNAbs have been isolated, few display pan-neutralizing activity and this subset is of particular value for clinical application or vaccine design^32–34,36^. This highlights the importance of comprehensive strategies to identify bNAbs that comprise critical characteristics required for effective clinical use. In this study, we thoroughly evaluated a carefully preselected cohort of HIV-1 elite neutralizers, including multiple donors with prolonged off-ART periods^65^, and performed detailed memory B cell analyses including functional characterization of 831 mAbs. Remarkably, only one of these antibodies—04_A06— emerged as a promising candidate for immunotherapy and prevention strategies underscoring the rarity of such findings and challenges that active vaccination efforts may have to overcome.

HIV-1 Env diversity and the rapid emergence of viral escape continue to represent a substantial challenge for successful clinical implementation of anti-HIV-1 bNAbs^2–5,8^. Therefore, achieving sufficient breadth and potency, along with prevention of viral escape, is crucial for the applicability of bNAbs. The antiviral activity of bNAbs is evaluated against large reference multiclade pseudovirus panels in order to compare neutralization profiles and to assess clinical applicability in a standardized manner^64,75,76^. However, these reference panels were generated based on virus strains circulating between 1998 and 2010 and may not reflect the current diversity and resistance of circulating HIV-1 strains. Indeed, the continuous evolution of HIV-1 led to a doubling of the concentration of VRC01 required to inhibit 50% of clade C viruses over the past 13 years^111^. HIV-1 strains isolated from the AMP trials represent a panel of contemporaneous-circulating HIV-1 strains and offer unique opportunities to evaluate the clinical potential of bNAbs against recently-transmitted viruses^9,102,112^. Notably, 04_A06 shows high activity against both traditional panels and viruses from the recent AMP studies. In addition to breadth and potency, restriction of viral escape is critical for maintaining antiviral activity in a clinical setting. By *in vitro* neutralization assays, deep mutational scanning analyses and *in vivo* treatment experiments, we demonstrated that 04_A06 effectively limits HIV-1 escape. Previous DMS studies of VRC01-class antibodies revealed distinct escape pathways involving canonical sites in loop D (e.g., N279_gp120_, A281_gp120_) and glycan-associated mutations near N197_gp120_, as well as distal sites like I326_gp120_^113^. Our results are consistent with these findings, as we also observed escape from VRC01-class antibodies N6LS, VRC07-523-LS and 3BNC117 via mutations at N279_gp120_ and/or A281_gp120_, and decreased neutralization sensitivity of 1-18 to mutations near the N197_gp120_ glycosylation motif. 04_A06 differed markedly from these antibodies in that no single mutation in our DMS conferred strong escape. However, our DMS analyses were conducted in the context of a single HIV-1 pseudovirus strain (HIV-1_BF520_) and the findings identified here may not be generalizable across other genetically diverse HIV-1 strains. Furthermore, the use of pseudoviruses is limited to capture the full spectrum of replication dynamics and evolutionary pressure present in replication-competent viral swarms. Nonetheless, in HIV-1_YU2_–infected humanized mice, 04_A06 achieved complete and sustained viral suppression extending 28 days beyond interruption of the treatment. Importantly, 04_A06 could overcome VRC01-class-resistant viruses circulating in pretreated mice and fully suppress viremia in all animals. These characteristics are uncommon among anti-HIV-1 bNAbs. In comparison to the V_H_1-46-encoded bNAb 1-18 which demonstrated prolonged suppression of viremia *in vivo*, 04_A06 is characterized by superior activity against CD4bs bNAb resistant virus strains and a unique escape profile identified through DMS^36^. The high antiviral activity and restriction of viral escape suggest that 04_A06 imposes strong selection pressure imposing fitness costs on HIV-1. While these findings highlight the clinical potential of 04_A06, our i*n vivo* studies were conducted using a single HIV-1 virus strain (YU2) and are limited to fully recapitulate viral diversity encountered in natural infection. Future studies incorporating a broader panel of viral strains or heterogeneous viral swarms could provide a more comprehensive assessment of potential escape pathways. This an important consideration for the clinical use of bNAbs against the diverse HIV-1 strains that circulate globally.

Structural analysis of 04_A06 revealed mechanisms that contribute to the antibody’s breadth and potency. Most notably, 04_A06 employs an ultra-long 11-amino acid insertion in FWRH1 that contacts K207_gp120_, H66_gp120_, and H72_gp120_, highly conserved residues on the adjacent gp120. Pseudoviruses containing substitutions at these residues exhibit decreased or completely abrogated infectivity^86^, suggesting these residues are functionally important and that escape mutations at these sites are likely associated with fitness costs. Consistently, engrafting the FWRH1 insertion from 04_A06 into VRC07 restored neutralizing activity against VRC01-class resistant viruses *in vitro*, further supporting the functional relevance of these contacts for restriction of viral escape. To our knowledge, 04_A06 is the first antibody that contacts this conserved surface on HIV-1 Env, a region that is sterically difficult for antibody access. In addition, an unliganded 04_A06 Fab structure showed that its FWRH1 insertion is pre-organized for binding, despite potential flexibility arising from it extending away from the antibody combining site, suggesting no entropic penalties for reorganizing the FWRH1 insertion upon Env binding. Beyond these heavy chain features, 04_A06 diverges from canonical VH_1-2_-enoded CD4bs bNAbs in its light chain architecture. Unlike VRC01 or VRC07, which exhibit a shortened CDRL1 to accommodate the N276_gp120_ glycan, 04_A06 lacks such deletion, highlighting alternative structural solutions for effective CD4bs engagement. In addition to 04_A06, we identified and structurally characterized 01_D03 and 05_B08, two phylogenetically distinct VRC01-class bNAbs also isolated from individual EN02. Like 04_A06, 01_D03 contacts the adjacent protomer, a likely consequence of its FWRH1 and FWRH3 insertions. The four-residue DASG FWRH3 insertion in 01_D03 is situated in the same position as a WDFD insertion previously identified in the 3BNC60/3BNC117 bNAb family, which arose in a different individual^37,114^. An analysis of clonal relatives of 3BNC60/3BNC117 revealed a correlation between the presence of this insertion and potent neutralizing activity, and its removal from 3BNC60 reduced its ability to neutralize diverse viruses^114^, further highlighting the potential importance of antibody insertions for breadth and potency.

Effective HIV-1 prevention approaches will be required to end the pandemic. Although promising concepts to guide B cell lineage development towards nAb responses are being explored^115–120^, an effective vaccine remains elusive. Passive immunization with bNAbs represents an alternative to active vaccination comparable to the use of long-acting antiretrovirals^9,121^. As revealed by the recent AMP trials, effective bNAb-mediated prevention strategies will critically depend on the use of bNAbs with both high potency and breadth^9,102^. Modelling indicates, that a single administration of 04_A06LS at standard dose and under standard assumptions of anti-HIV-1 bNAb *in vivo* half-life provides similar prevention efficacy as triple bNAb combinations that are being investigated in early-phase clinical trials^1,102^. However, these *in silico* predictions are based on pharmacokinetic data from the wild-type and LS-modified antibody in hFcRn mice. Therefore, this approach depends on assumptions that may not accurately reflect pharmacokinetics in humans. Among groups at risk of HIV-1 infection, newborn children remain particularly vulnerable^122^. Although, antiretrovirals have been authorized for postnatal prophylaxis, drug tolerance and dose finding in children remains challenging^123–125^. Especially in these groups, passive administration of long-acting bNAbs is attractive due to favorable safety profiles, extended protection provided over the duration of breastfeeding^126^ and high treatment adherence^127^. Therefore, bNAbs such as 04_A06 are essential to ensure antibody-mediated prevention to be an effective and cost-saving solution for HIV-1 prevention.

In summary, by conducting the most extensive single cell analysis to date on a cohort of top HIV-1 elite neutralizers, we identified CD4bs bNAb 04_A06 that achieves high antiviral activity and restriction of viral escape through unique structural features and has promising potential for clinical development.

## Methods

### Study participants and collection of clinical samples

Large-blood draw and leukapheresis samples were collected according to protocols reviewed and approved by the Institutional Review Board (IRB) of the University of Cologne (study protocols 13-364 and 16-054) and local IRBs. Study participants were recruited at private practices and/or hospitals in Germany (Cologne, Essen, and Frankfurt), Cameroon (Yaoundé), Nepal (Kathmandu), and Tanzania (Mbeya) and provided written informed consent. In total, serum samples of 2,354 study participants were screened for anti-HIV-1 neutralizing activity in order to identify HIV-1 elite neutralizers^65^. Subsequently, 32 elite neutralizers among the top 3.7% of the cohort were selected for large blood draw collection and B cell isolation. Clinical information was extracted from medical records.

### Cell lines

HEK293T cells were maintained in Dulbecco’s Modified Eagle Medium (DMEM, Thermo Fisher) supplemented with 10% fetal bovine serum (FBS, Sigma-Aldrich), 1x antibiotic-antimycotic (Thermo Fisher), 1 mM sodium pyruvate (Gibco) and 2 mM L-glutamine (Gibco) at 37 °C and 5% CO_2_. HEK293-6E cells were maintained in FreeStyle 293 Expression Medium (Life Technologies) supplemented with 0.2% penicillin/streptomycin under constant shaking at 90 – 120 rpm, 37 °C and 6% CO_2_. TZM-bl cells (Platt et al., 1998) were cultured at 37 °C in 5% CO2 i,n DMEM supplemented with 10% FBS, 1 mM sodium pyruvate, 2 mM L-glutamine, 50 µg/ml gentamicin (Merck), and 25 mM HEPES (Millipore). The sex of HEK293T, HEK293-6E and TZM-bl cell lines is female. Cell lines were not specifically authenticated.

### Mouse Models

NOD.Cg-Rag^1tm1mom^Il2rg^tm1Wjl^/SzJ (NRG) mice obtained from The Jackson Laboratory and subsequently bred and housed at the University of Cologne’s Decentralized Animal Husbandry Network (Dezentrales Tierhaltungsnetzwerk) of the University of Cologne. These mice were kept in specific pathogen-free (SPF) conditions with a 12-hour light/dark cycle. For breeding, the mice received ssniff 1124 breeding feed, whereas those used for experimental purposes were given ssniff 1543 maintenance feed. The creation of humanized mice followed a previously-established method, with some modifications^8,129^. Human CD34+ hematopoietic stem cells were harvested from cord blood and placental tissue using CD34 microbeads for immunomagnetic cell separation (Miltenyi Biotec). Both cord blood and placental tissues were obtained with written informed consent under a protocol approved by the University of Cologne’s Institutional Review Board (16-110) or Ethics Committee of the Medical Association of North Rhine (2018382). Within five days of birth, NRG mice received sublethal irradiation and were injected intrahepatically with the isolated human CD34+ stem cells 4 to 6 hours later. Twelve weeks post-injection, successful humanization was confirmed through FACS analysis of mouse blood for human PBMCs^8^. Already humanized NOD-Prkdc^scid^-IL2rg^Tm1^/Rj (NXG-HIS) mice were used as a complementary model and were directly obtained from Janvier Labs. All procedures involving these mice were sanctioned by the State Agency for Nature, Environmental Protection, and Consumer Protection North Rhine-Westphalia (LANUV).

### Isolation of PBMCs and plasma

Peripheral blood mononuclear cells (PBMCs) were isolated from large blood draws of leukapheresis by density gradient separation using Histopaque separation medium (Sigma-Aldrich) and Leucosep cell tubes (Greiner Bio-one) according to the manufacturer’s instructions. Isolated PBMCS were stored at −150 °C in 90% FBS supplemented with 10% DMSO until further use. Plasma was collected and stored at −80 °C.

### Isolation of single anti-HIV-1-reactive B cells

The isolation of single antigen-reactive B cells was performed as previously described^63,130^. In brief, fractions of CD19^+^ B cells were enriched from PBMCs using immunomagnetic cell separation with CD19 microbeads (Miltenyi Biotec) according to the manufacturer’s protocol. Subsequently, enriched CD19+ B cells were stained with 4’,6-Diamidin-2-phenlindol (DAPI; Thermo Fisher Scientific), anti-human CD20-Alexa Fluor 700 (BD), anti-human IgG-APC (BD), and the GFP-labeled BG505_SOSIP.664_ or biotinylated Streptavidin-labeled YU2gp_140_ HIV-1 Env bait (15 µg/ml) on ice for 20 min. DAPI^-^ CD20^+^ HIV-1 Env-reactive IgG single cells were sorted into 96 well plates applying a FACSAria Fusion (Becton Dickinson). Each well of the 96 well plate was prefilled with 4 µl of sorting buffer consisting of 0.5 x PBS, 0.5 U/µl RNAsin (Promega), 0.5 U/µl RNaseOut (Thermo Fisher Scientific), and 10 mM DTT (Thermo Fisher Scientific). Plates were cryopreserved at −80 °C immediately after sorting.

### B cell receptor amplification and sequence analysis

Generation of cDNA and amplification of antibody heavy and light chain genes from sorted single cells was performed as previously described^36,63,131^. For reverse transcription, isolated HIV-1-reactive B cells were incubated with 7 µl of a random-hexamer-primer master mix (RHP mix) consisting of 0.75 µl Random Hexamer Primer (Thermo Fisher Scientific), 0.5 µl NP-40 (Thermo Fisher Scientific), 0.15 µl RNaseOut (Thermo Fisher Scientific), and 5.6 µl of nuclease-free H_2_O at 65 °C for 1 min. Subsequently, reactions were incubated with 3 µl of 5 x Superscript IV RT buffer (Thermofisher Scientific), 0.5 µl dNTPs (Thermo Fisher Scientific), 1 µl DTT (Thermo Fisher Scientific), 0.1 µl of RNasin (Promega), 0.1 µl of RNaseOut (Thermo Fisher Scientific), 2.05 µl of nuclease-free H_2_O and 0.25 µl of Superscript IV (Thermofisher Scientific) per well at 42 °C for 10 min, 25 °C for 10 min, 50°C for 10 min and 94 °C for 5 min. Heavy and light chains of the B cell receptors were amplified from generated cDNA using semi-nested single cell PCRs using optimized V gene-specific primer mixes^68^ and Platinum Taq DNA Polymerase or Platinum Taq Green Hot Start Polymerase (Thermo Fisher Scientific) as previously described^131,36,68,63^. Amplicons were analyzed by agarose gel electrophoresis for correct PCR product size and subsequently shipped for Sanger sequencing. Only chromatograms with a mean Phred score of 28 and sequences with a minimal length of 240 nucleotides were selected for downstream sequence analyses. Filtered sequences were annotated with IgBlast^132^ according to the IMGT system^133^ and only the variable region from FWR1 to the end of the J gene was extracted. Base calls within the variable region with a Phred score below 16 were masked and sequences with more than 15 masked nucleotides, stop codons, or frameshifts were excluded from further analyses. Sequence analyses to inform on sequence clonality were performed separately for each study participant. All productive heavy chain sequences were grouped by identical V_H_/J_H_ gene pairs and the pairwise Levenshtein distance for their CDRH3s was determined. Starting from a random sequence, clone groups were assigned to sequences with a minimal CDRH3 amino acid identity of at least 75% (with respect to the shortest CDRH3). 100 rounds of input sequence randomization and clonal assignment were performed and the result with the lowest number of remaining unassigned (non-clonal) sequences was selected for downstream analyses. All clones were cross-validated by the investigators taking shared mutations and light chain information into account.

### Cloning and production of mAbs

Heavy and light chain variable regions of selected antibodies were cloned into expression vectors using sequence- and ligation-independent cloning (SLIC) as previously described^134,36,131,63,130^. The product of the 1^st^ PCR was amplified using Q5 Hot Start High Fidelity DNA Polymerase (New England Biolabs) and specific forward and reverse primers including adaptor sequences which are homologous to the restriction sites of the antibody expression vector (IgG1, IgL, IgK)^134^. Multiplexed forward primers were designed according to 2^nd^ PCR primers and encode for the complete native leader sequence of all heavy and light chain V genes, whereas reverse primers bind to the conserved sequence motifs at the 5’ end of heavy and light chain immunoglobulin constant regions^63,68^. Reactions were incubated at 98 °C for 30 s; 35 cycles of 98 °C for 10 s, 72 °C for 45 s; and 72 °C for 2 min and PCR products were purified using 96-well format silica membranes. Modified antibody versions in which amino acid insertions were added or deleted were generated by cloning of a synthesized heavy chain variable region DNA fragment including the insertions or deletions (Integrated DNA Technologies). Purified PCR products and/or synthesized heavy chain variable region DNA fragments were cloned into linearized expression vectors with T4 DNA polymerase (NEB) and transformed into chemically competent *Escherichia coli* (DH5α). Correct insertion of the V region sequences into the expression vectors was confirmed by Sanger sequencing and positive colonies were further propagated. Subsequent experimental procedures were performed either as a high-or low-throughput approach as previously described^63^. The high-throughput approach was applied antibody production and screening with the purpose to rapidly identify neutralizing antibodies from 32 HIV-1 elite neutralizers. Screening hits were then produced following the low-throughput approach that yields antibody quantities sufficient for more resource-consuming experiments. Monoclonal antibodies were recombinantly produced in HEK293-6E suspension cells or HEK293T adherent cells by co-transfecting human heavy chain and corresponding light chain antibody expression vectors using 25 kDa branched polyethylenimine (PEI; Sigma-Aldrich) or TurboFect Transfection Reagent (Thermo Fisher Scientific). Transfected HEK293-6E cells were propagated in FreeStyle 293 Expression Medium (Thermo Fisher Scientific) supplemented with 0.2% penicillin/streptomycin (Thermo Fisher Scientific) at 37 °C and 6% CO_2_ under constant shaking at 90 – 120 rpm for five to seven days. Transfected adherent HEK293T cells were maintained in Dulbecco’s Modified Eagle Medium (DMEM, Thermo Fisher) supplemented with 10% fetal bovine serum (FBS, Sigma-Aldrich), 1 x antibiotic-antimycotic (Thermo Fisher), 1 mM sodium pyruvate (Gibco) and 2 mM L-glutamine (Gibco) for four days.

To produce Fabs for cryo-EM analysis, heavy chain variable regions of 04_A06, 05_B08, and 01_D03 were subcloned into a p3BNC-based Fab expression vector encoding a C-terminal hexahistidine tag and expressed with corresponding light chain-encoding plasmids via transient co-transfection of Expi293F cells (Thermo Fisher). Fabs were purified from culture supernatant by Ni-NTA affinity chromatography (GE Life Sciences). Fabs were concentrated and buffer-exchanged into TBS (20mM Tris pH 8.0, 150mM NaCl) using Amicon 10 kDa spin concentrators (Millipore) and further purified by size-exclusion chromatography (SEC) on a Superdex-200 16/60 column equilibrated with TBS. SEC fractions corresponding to the Fab peak were either directly used or combined and concentrated using an Amicon 10 kDa spin concentrator (Millipore) and stored at 4°C.

### Expression and purification of BG505 SOSIP trimer

BG505_SOSIP.664_ used for cryo-EM was expressed via transient co-transfection with a furin-encoding expression plasmid in Expi293F cells (Thermo Fischer) as described^135^. SOSIP was purified from cell culture supernatants by either 2G12 (04_A06/PGDM1400 complex) or PGT145 (01_D03 and 05_B08 complexes and unbound structure) immunoaffinity chromatography, dialyzed into TBS, and concentrated. SEC was performed on a Superose-6 Increase column (PGT145-purified prep) or a Superdex-200 16/60 column followed by a Superose-6 Increase 10/300 GL column (2G12-purified prep) equilibrated in TBS. SEC fractions were stored individually at 4°C prior to use.

### Protein G-based antibody purification

Cell supernatants from HEK293-6E suspension cells, which were transiently transfected with heavy and light chain expression vectors, were collected by centrifugation, filtered through PES filters (Cytiva), and incubated overnight at 4°C with Protein G-coupled Sepharose beads (GE Life Sciences). The suspension was transferred to chromatography columns, rinsed twice with phosphate-buffered saline (PBS), and IgGs were eluted with 0.1 M glycine (pH 3) and buffered using 1 M Tris (pH 8). Subsequently, buffer exchange to PBS and antibody concentration were performed by centrifugation using 30 kDa Amicon spin membranes (Millipore). Purified were filtered through Ultrafree-MC 0.22 µm membranes (Millipore) and stored at 4 °C until further use. The final concentration of purified antibodies was measured by UV/Vis spectroscopy using a Nanodrop (A_280_ nm).

### Purification of serum IgGs

Serum was diluted 1:1 with Dulbecco’s PBS (DPBS; Gibco, Thermo Fisher Scientific). IgG was isolated by passing the diluted serum three times through a column containing 1.5 ml of protein G agarose (Pierce, Thermo Fisher Scientific). The column was then washed with DPBS and the bound IgG was eluted using glycine-HCl (pH 2.7). The eluate was immediately neutralized with 1 M Tris-HCl (pH 8.0). To concentrate the IgG and remove salts, the buffer was exchanged to PBS using a 15 ml tube with a 10 kDa molecular weight cut-off (MWCO) filter (Thermo Fisher Scientific).

### Determination of antibody concentrations by human IgG capture ELISA

The human IgG capture ELISA was applied to determine antibody concentrations in unpurified cell supernatants of transfected HEK293T or HEK293-6E cells as previously described with minor modifications^63^. In brief, ELISA plates (Greiner Bio-One) were coated with 2.5 µg/ml of polyclonal goat-anti-human IgG in PBS at least for 45 min at 37°C or at 4 °C overnight. Plates were blocked with blocking buffer (BB) consisting of PBS supplemented with 5% nonfat dried milk powder (Carl Roth T145.2) for 60 min at RT. Cell supernatants of transfected HEK293T cells were tested with a starting dilution of 1:20 in BB. Human myeloma IgG1 kappa from human myeloma plasma (Sigma Aldrich) was used at a starting concentration of 4 µg/ml in BB as a standard. Subsequently samples were serially diluted 1:3 in BB. After incubation for 45 min at RT, plates were incubated with anti-human IgG-HRP (Southern Biotech 2040-05) diluted 1:2500 in BB. Plates were developed using ABTS solution (Thermo Fisher Scientific 002024) and absorbance was measured at 415 nm and 695 nm by a Microplate Reader (Tecan). Antibody concentration in cell culture supernatants was determined by comparison with the human myeloma IgG1 standard.

### HIV-1 pseudovirus production

HIV-1 pseudoviruses were produced in HEK293T cells by co-transfection with pSG3Δenv and the respective HIV-1 Env plasmids as previously described^69,71,74,76^. For donor- and mouse-derived sequences, HIV-1 Env plasmids were synthesized by Twist Bioscience.

### Neutralization assay

Neutralization assays were performed to determine the IC_50/80_ of purified mAbs and purified serum IgGs for donor screening and/or to screen cell culture supernatants of transfected HEK293T cells without IgG purification for neutralizing activity. Neutralization assays were conducted as previously described with minor modifications^69^. For the large-scale donor screening, isolated IgGs from participants were tested against each virus in a single well at a concentration of 300 µg/ml in duplicates. Monoclonal antibodies, purified serum IgGs or cell culture supernatants of transfected HEK293T cells were incubated with pseudovirus strains for 1h at 37 °C, followed by the addition of TZM-bl cells at a final concentration of 10^4^ cells per well in a 96-well plate. After a 48h incubation period at 37 °C and 5% CO_2_, luciferase activity was assessed applying the luciferin/lysis buffer. After subtracting background RLUs of non-infected cells, % of neutralization was calculated and used for reporting. For testing of unpurified IgGs derived from HEK293T cell culture supernatants, a final concentration of 2.5 µg/ml was used. The concentration of IgG in cell culture supernatants was determined by human IgG capture ELISA. To determine IC_50/80_ values for mAbs, a dilution series of the mAb was performed starting with a concentration of 10, 25, or 50 µg/ml. IC_50/80_ values were calculated as the mAb concentration causing a 50% or 80% reduction in signal compared to the virus-only controls using a dose-response curve in GraphPad Prism, respectively. For the determination of IC_50/80_ values, each mAb was tested in duplicates. Produced reference antibodies were functionally validated through neutralization assays against the global HIV-1 pseudovirus panel. The resulting IC₅₀ and IC₈₀ values for each antibody were compared to historical data available in the CATNAP database^128^. Only those antibodies exhibiting less than a threefold deviation in IC₅₀/IC₈₀ values relative to the reference data were included in subsequent functional analyses.

### Antibody Epitope Prediction Using Neutralization Fingerprinting

Epitope prediction of serum IgG neutralizing activity was performed using a computational approach as previously described^71,136^. Neutralization was measured in a TZM-bl assay against a panel of 20 HIV-1 pseudoviruses comprising the f61 fingerprinting panel^71^. The resulting polyclonal neutralization profiles were deconvoluted into contributions from ten epitope-specific antibody clusters, previously defined by clustering broadly neutralizing reference antibodies according to their neutralization fingerprints^136^. The prevalence of each epitope specificity within a given serum was estimated via a least-squares fitting procedure, yielding a vector of ten delineation scores—one per antibody cluster—each ranging from 0 (low prevalence) to 1 (high prevalence) and collectively summing to one. A threshold of 0.25 was applied to classify a score as a positive signal, thereby allowing identification of up to four dominant specificities per serum.

### HIV-1 Env ELISAs

High-binding ELISA plates (Greiner Bio-One) were coated an HIV-1 Env protein at a concentration of 4 µg/ml. Plates were blocked with 2% BSA (Sigma Aldrich) and 0.05% Tween-20 (Carl Roth) in PBS for 1h at 37°C. Monoclonal antibodies were tested at a starting concentration of 10 µg/ml in PBS and were serially diluted 1:4. After incubation with antibodies for 1h at RT, plates were incubated with anti-human IgG-HRP (Southern Biotech 2040-05) diluted 1:1,000 in 2% BSA in PBS for 1h at RT. Between each step, plates were washed with PBS containing 0.05% Tween-20 (Carl Roth). All samples were tested in duplicates. Plates were developed using ABTS solution (Thermo Fisher Scientific 002024) and absorbance was measured at 415 nm and 695 nm by a Microplate Reader (Tecan).

### Competition ELISAs

Selected mAbs were biotinylated using the EZ-Link Sulfo-NHS-Biotin Kit (Thermo Fisher Scientific) according to the manufacturer’s protocol. Buffer exchange to PBS was performed by centrifugation using Amicon 10 kDA centrifugation filter membranes (Millipore). High-binding ELISA plates (Greiner Bio-One) were coated with anti-6x His tag antibody (Abcam 9108) at a concentration of 2 µg/ml overnight at 4°C. Plates were blocked with 3% BSA (Sigma Aldrich) in PBS for 1h at 37°C. After blocking, wells were incubated with BG505_SOSIP.664_-His at a concentration of 2 µg/ml in PBS for 1h at RT. Competing antibodies were used at a starting concentration of 32 µg/ml in PBS and serially diluted 1:3. After incubation with competing antibodies for 1h at RT, biotinylated antibodies of interest were added at a concentration of 0.5 µg/ml in PBS containing 3% BSA and plates were incubated for 1h at RT. Subsequently, plates were incubated with peroxidase-streptavidin (Jackson ImmunoResearch) diluted 1:5,000 in PBS containing 1% BSA and 0.05% Tween-20. Between each step, plates were washed with PBS containing 0.05% Tween-20 (Carl Roth). Plates were developed using ABTS solution (Thermo Fisher Scientific 002024) and absorbance was measured at 415 nm and 695 nm by a Microplate Reader (Tecan).

### Autoreactivity evaluations in HEp-2 cell assays

HEp-2 cell autoreactivity was analyzed using the NOVA Lite Hep-2 ANA Kit (Inova Diagnostics) following the manufacturer’s protocol. MAbs were utilized at a concentration of 100 μg/ml in PBS.

Images were captured with a DMI 6000 B fluorescence microscope (Leica) set to a 3-second exposure time, 100% intensity, and a gain of 10.

### Cryo-EM sample preparation

An 04_A06-PGDM1400-BG505 complex was set up in a 3.6:1.2:1 molar ratio of 04_A06 Fab: PGDM1400 Fab: BG505 trimer and incubated at RT overnight. Complex was purified on a Superose-6 Increase 10/300 GL column and SEC fractions corresponding to the leading half of the complex peak were pooled and concentrated using an Amicon 10 kDa spin concentrator (Millipore) to ∼2.5 mg/ml in TBS one day prior to vitrification. For the 01_D03-BG505 and 05_B08-BG505 complexes, Fabs were incubated overnight in TBS at room temperature in a 3.6:1 molar ratio of Fab: BG505 trimer. Complexes were concentrated to ∼4.2 mg/ml (01_D03 complex) or ∼4.4 mg/ml (05_B08 complex) using an Amicon 10 kDa spin concentrator (Millipore) without additional SEC purification. The unbound BG505 sample was concentrated to ∼4.0 mg/ml.

Octyl-maltoside, fluorinated solution was added to each sample for a final concentration of 0.02% (w/v) immediately preceding addition of 3 µL to a Quantifoil R1.2/1.3 Cu 300 mesh grid (04_A06/PGDM1400 complex) or a Quantifoil R1.2/1.3 Holey Carbon Film, 300 mesh gold grid (01_D03 and 05_B08 complexes and unbound BG505; Electron Microscopy Services) that had been glow discharged for 1 min at 20 mA using a PELCO easiGLOW (Ted Pella). Grids were blotted for 3s with Whatman No. 1 filter paper and plunge frozen in liquid ethane using a Mark IV Vitrobot (Thermo Fisher) operating at RT and 100% humidity.

### Cryo-EM data collection and processing

Data were collected on a 300 keV Titan Krios transmission electron microscope (Thermo Fisher Scientific) equipped with a GIF Quantum energy filter and a K3 6k x 4k direct electron detector (Gatan) operating in counting mode. Data collection was performed using SerialEM v4.0.13 (04_A06 and unbound BG505) or v4.1.0beta^137^ (01_D03 and 05_B08) at a nominal magnification of 105,000x (super-resolution = 0.416Å/pixel) and a defocus range of −1.0 to −3.0 µm. Movies were recorded using a 3x3 beam image shift pattern with 1 shot (04_A06/PGDM1400 dataset) or 2 shots (01_D03 and 05_B08 datasets), or 3 shots (unbound BG505 dataset) per hole. For the 04_A06 and 01_D03 datasets, motion correction, CTF estimation, particle picking, and binned particle extraction were performed using cryoSPARC Live (v3 and v4, respectively) before processing in cryoSPARC^137^. For the 05_B08 and unbound BG505 datasets, all processing was carried out in cryoSPARCv4. Particles were picked using blob picker or Topaz^138^ and extracted from micrographs. Further processing details for individual datasets can be found in Extended Data Fig. 5 and Supplementary Table 9. For the 04_A06-PGDM1400-BG505 dataset, a model was built into the map containing PGDM1400, and after verifying that this model (minus PGDM1400 and the N160 glycans) fit well into the C3 symmetric map that lacked PGDM1400, particle subtraction and local refinement (with applied C3 symmetry) was carried out to obtain a higher resolution view of the 04_A06-BG505 interface. To create a mask for particle subtraction, the “molmap” command in ChimeraX was applied to a model of the PGDM1400 Fab and N160_gp120_ glycans that was built into the EM density. The mask was imported into cryoSPARC^137^, and a soft padding applied (threshold = 0.1; soft padding width = 10 voxels). A mask for local refinement was similarly created using a model of BG505 and the 04_A06 VHVL (threshold = 0.05, dilation radius = 5; soft padding width = 10).

### Structure modelling and refinement

The following coordinates were docked into the corresponding densities of the EM maps using ChimeraX^91^ to generate starting models: BG505 (PDB 6UDJ)^36^, PGDM1400 (PDB 4RQQ)^79^, and the 04_A06 Fab crystal structure (PDB 8UKI; this study). Sequence-corrected models for the 01_D03 and 05_B08 VHVL domains were built into the corresponding densities in Coot^139^ using the 04_A06 Fab crystal structure as a starting model. Models were refined through iterative rounds of Phenix real space refine and Coot. N-glycans were built using tools in Coot^139^ and verified as ‘OK’ by Privateer^140^. Modelling of side chains should be considered approximate, owing to the intermediate resolution of the EM structures. Antibody residues were numbered according to Kabat.

### X-ray crystallography

Crystallization screens for 04_A06 Fab were performed using sitting drop vapor diffusion at RT by mixing 0.2 µL Fab (4.1 mg/ml) with 0.2 µL of reservoir solution (Hampton Research) using a TTP Labtech Mosquito automatic microliter pipetting robot. 04_A06 Fab crystals were obtained in 8% v/v TacsimateTM pH 7.0, 20% w/v polyethylene glycol 3,350. Crystals were looped and cryopreserved in reservoir solution supplemented stepwise with 5-20% glycerol and cryopreserved in liquid nitrogen. A 1.75 Å structure of 04_A06 Fab was solved using a data set collected at 100 K and 1 Å wavelength on Beamline 12-2 at the Stanford Synchrotron Radiation Lightsource (SSRL) with an Eiger X 16M (Dectris) detector, which was indexed and integrated with iMosflm v7.4^141^ and then merged with AIMLESS in the CCP4 software package v7.1.018^80^.

The structure was determined by molecular replacement in Phaser with the coordinates of the VRC01 Fab (PDB 3NGB), using the CH-CL and VH-VL (with truncated CDR loops) as separate search models. Coordinates were refined using PHENIX v1.20.1-4487129^142^ with individual B factors and TLS restraints130^143^. Manual rebuilding was performed iteratively with Coot v0.9.8.8131^144^. >98% of residues were in the favored regions of the Ramachandran plot and <1% in the disallowed region (Supplementary Table 9).

### Structural analyses

Figures were prepared using UCSF ChimeraX^90,91^ and PyMOL (Schrödinger LCC). Buried surface area was calculated using PDBePISA v1.52^145^ with a 1.4Å probe. RMSDs were calculated in PyMOL (Schrödinger LCC) and electrostatic surfaces were calculated in UCSF ChimeraX^90,91^ . Owing to the intermediate resolution and minor differences in modelling of identical copies of chains within a trimer, a general cut-off of ::6.0Å was used to define potential interactions.

### Generation of mutant HIV-1_YU2_ and HIV-1_BG505_ pseudovirus mutants

Point mutations were introduced into HIV-1_YU2_ or HIV-1_BG505_ gp160 expression plasmids using the Q5 Site-Directed Mutagenesis Kit (New England Biolabs) according to the manufacturer’s protocol. Pseudovirus mutants were produced as described above.

### Deep mutational scanning

A lentivirus deep mutational scanning platform was applied to quantitatively assess the impact of mutations and combinations thereof on the functionality of the HIV-1 env protein and their resistance to neutralization by antibodies as previously described^98,99^. For the present investigation, we deployed mutant libraries of the HIV-1 virus strain BF520 to explore antibody resistance patterns. In brief, two distinct BF520 env mutant lentivirus libraries were used, each containing around 40,000 mutants with an average of approximately 2.5 nonsynonymous mutations per mutant. For antibody selections, VSV-G pseudotyped neutralization standard viruses were added to virus pools to make up 0.5-1% of the total. Each selection condition involved incubating 1 million infectious units from one of the mutant libraries containing the pseudotyped standard with various dilutions of antibodies, spanning concentrations from IC90 to IC99.9 along with a mock incubation for control purposes. Following a one-hour incubation period, the mixtures were used to infect 1 million TZM-bl cells per well in a six-well plate, with the addition of 100 µg/mL DEAE dextran to enhance infection efficiency. Twelve hours post-infection, the unintegrated lentivirus genomes from each condition were extracted and was processed for sequencing as previously described^99^.

### Deep mutational scanning data analysis

The dms-vep-pipeline-3 (version 3.2.3) was applied to analyze generated deep mutational scanning data (https://github.com/dms-vep/dms-vep-pipeline-3) and the full analysis including CSV files with the numerical measurements and HTML renderings of key analyses was deposited in a GitHub repository (https://dms-vep.github.io/HIV_Envelope_BF520_DMS_04-A06). Analysis of PacBio and Illumina sequencing data, as well as modeling of mutation effects on HIV Envelope function and escape from neutralizing antibodies, were conducted as previously described^98,99^. In brief, to model antibody escape, we estimated the fraction of each mutant that was not neutralized in each antibody selection by comparing the counts of each mutant to those of the non-neutralized standard viruses under both antibody and mock incubation conditions. In order to quantify the effects of each individual mutation on escape from each antibody, the software package polyclonal^100^ version 6.6 (https://jbloomlab.github.io/polyclonal) was applied. The logo plots (Extended Data Fig. 8) display the modeled effects of each individual mutation on escape from each antibody, where the height of the letter of the amino-acid represents the magnitude of the measured effect on escape.

### Production of recombinant HIV-1 virus

Replication-competent recombinant HIV-1 (YU2 *env* in NL4-3 backbone^146^) was produced by transfection of HEK293T cells using FuGENE 6 Transfection Reagent (Promega). Cell culture supernatants containing recombinant virus were harvested 48h or 72h after transfection and stored at −80°C.

### Viral outgrowth of replication-competent isolates

CD4+ T cells were isolated from the peripheral blood mononuclear cells (PBMCs) of PLWH using the CD4+ T Cell Isolation Kit from Miltenyi Biotec. These cells were then co-cultured with irradiated (50 Gy) PBMCs from healthy donors in T cell medium, which consisted of RPMI 1640 supplemented with 300 mg/l L-glutamine (Thermo Fisher), 10% FBS (Sigma-Aldrich), and 1% penicillin/streptomycin (Thermo Fisher). The cultures were treated with 1 μg/ml PHA-M (Sigma-Aldrich) and 100 U/ml interleukin-2 (IL-2) (Miltenyi Biotec). After one day, the medium was replaced with fresh T cell medium containing 100 U/ml IL-2 and 5 μg/ml polybrene (Sigma-Aldrich). Additionally, stimulated healthy donor PBMCs, which had been treated for two days with 1 μg/ml PHA-M and 100 U/ml IL-2, were added to the culture. Prior to their addition, the donor PBMCs were depleted of CD8+ T cells using CD8 MACS MicroBeads (Miltenyi Biotec). Weekly supplementation of the culture with additional CD8+ T cell-depleted donor PBMCs was performed. The culture supernatants were regularly monitored for p24 antigen production using the Architect HIV Ag/Ab Combo Assay (Abbott). Once p24 was detected, the supernatants were harvested and stored at temperatures ranging from −80°C to −150°C.

### Infection of humanized mice and viral load measurements

Humanized NRG mice were challenged with replication-competent HIV-1_YU2_ intrapertioneally. 21-24 days after viral challenge, infected mice were treated subcutaneously by injection of sterile monoclonal antibodies in PBS. 1 mg of antibodies was applied as a loading dose, followed by injections of 0.5 mg every 3 – 4 days. For viral load measurements, viral RNA was isolated from EDTA plasma samples using the MinElute Virus Spin Kit (Qiagen) and DNase I (Qiagen) on an automated Qiacube (Qiagen). Viral loads were determined by quantitative real-time PCR (qRT-PCR) using *pol*-specific primers 5’TAATGGCAGCAATTTCACCA and 5’GAATGCCAAATTCCTGCTTGA, and 5’/56-FAM/CCCACCAACARGCRGCCTTAACTG/ZenDQ/ as probe, as previously described^50^. qRT-PCR was conducted on a QuantStudio 5 (Thermo Fisher Scientific) using the Taqman RNA-to-CT 1-Step-Kit (Thermo Fisher Scientific). As standard for the determination of viral loads, heat-inactivated culture supernatants of replication-competent HIV-1_YU2_ propagated by infection of SupT1-R5 cells were included in every PCR run. The viral load of the standard was assessed using the quantitative cobas 6800 HIV-1 kit (Roche). The lower limit of detection for the qRT-PCR was determined as 784 copies/ml. For calculations of log_10_ changes, viral loads below 784 copies/ml were assigned a copy concentration 783 copies/ml.

### Single genome sequencing of plasma HIV-1 env genes from humanized mice

Single Genome Sequencing (SGS) was performed as previously described^101^. In brief, viral RNA was isolated from EDTA plasma samples using the MinElute Virus Spin Kit (Qiagen) and DNase I (Qiagen) on an automated Qiacube (Qiagen). Reverse transcription was performed using the antisense primer YB383 5’TTTTTTTTTTTTTTTTTTTTTTTTRAAGCAC^147^ and SuperScript IV (Thermo Fisher Scientific) according to the manufacturer’s instructions and followed by an incubation with 0.25 U/µl RNase H (Thermo Fisher Scientific) at 37°C for 20 min. Subsequently, Env-encoding cDNA was serially diluted and amplified by a nested PCR using Platinum Taq Green Hot Start (Thermo Fisher Scientific) and HIV-1_YU2-NL4-3_ specific primers. For the 1^st^ PCR primers YB383 and YB50 5’ GGCTTAGGCATCTCCTATGGCAGGAAGAA were used and reactions were incubated at 94°C for 2 min; 35 cycles of 94°C for 30 sec, 55°C for 30 sec, and 72°C for 4 min; and 72°C for 15 min. Subsequently, 1 µl of 1^st^ PCR product was applied as a template for the 2^nd^ PCR. For the 2^nd^ PCR, primers YB49 5’ TAGAAAGAGCAGAAGACAGTGGCAATGA and YB52 5’ GGTGTGTAGTTCTGCCAATCAGGGAAGWAGCCTTGTG were used and reactions were incubated at 94°C for 2 min; 45 cycles of 94°C for 30 sec, 55°C for 30 sec, and 72°C for 4 min; and 72°C for 15 min. PCR products from amplifications < 30% efficiency were selected for purification using the NucleoSpin 96 PCR Clean-up Kit (Macherey-Nagel).

### Illumina dye sequencing of humanized mice SGS-derived *env* amplicons

SGS-derived *env* amplicons were purified with the NucleoSpin 96 PCR Clean-up Kit (Macherey-Nagel) and subjected to library preparation and NovaSeq 2x 150 bp sequencing at the Cologne Center for Genomics sequencing core facility.

### Determination of antibody PKs *in vivo*

For determination of wildtype antibody (04_A06, 10-1074, 3BNC117) half-life, human FcRn transgenic mice (The Jackson Laboratory) were administered 0.5 mg of purified antibody in PBS via intravenous tail vein injection. Total serum or plasma (bNAb-levels following treatment interruption) concentrations of human IgG were measured by ELISA, with minor modifications as previously described^8^. Briefly, high-binding ELISA plates (Corning) were coated overnight with anti-human IgG (Jackson ImmunoResearch) at 2.5 μg/ml at room temperature (RT). The wells were then blocked using a buffer containing 2% BSA (Carl Roth), 1 μM EDTA (Thermo Fisher), and 0.1% Tween 20 (Carl Roth) in PBS. A standard curve was generated using human IgG1 kappa purified from myeloma plasma (Sigma-Aldrich) diluted in PBS. Serial dilutions of the IgG standard and serum or plasma samples were incubated for 90 minutes at RT, followed by HRP-conjugated anti-human IgG (Jackson ImmunoResearch) at a 1:1,000 dilution in blocking buffer for another 90 minutes at RT. After adding ABTS (Thermo Fisher), the optical density at 415 nm was determined using a Tecan microplate reader. Each step included washing with 0.05% Tween 20 in PBS. Baseline serum or plasma samples confirmed the absence of human serum or plasma IgG before antibody injection.

For PK determination of 04_A06LS and Sotrovimab, human FcRn female mice (n=5) were weighed the day before injection and then received a single IV bolus dose of 5 mg/kg of test article via the tail vein, diluted in TA vehicle at 1 mg/ml and injected in a volume of 5 mL/kg. Blood was collected and processed to serum for pharmacokinetic testing at 15 timepoints from the time of infusion through 63 days post dose. Blood samples were collected from the mouse’s lateral tail veins via direct venipuncture. After collection, the tubes were left to sit in an upright position at room temperature for 30-45 min to allow complete clotting. The samples were centrifuged for 5 min at 2-8 °C, 5500 rpm and the resulting serum was frozen at −20°C until bioanalytical quantification. Sotrovimab, an anti-CoV2 mAb, was used as a comparator. A sandwich-based electrochemiluminescence ligand binding assay (MSD) was developed to measure the concentrations of the mAbs using an anti-LS mutation as capture antibody and an anti-huCH2 domain-sulfotag as detector. Pharmacokinetic parameters were estimated using WinNonlin (version 8.4.0.6172, Certara L.P., Princeton, NJ). The nominal dose levels and nominal sampling times were used in the calculation of all pharmacokinetic parameters. Standard noncompartmental methods and linear up/log down rules were used.

### Determination of the predicted PE and geometric mean PT_80_

The predicted PE and PT_80_ were determined following the methodology previously described^102^. Calculations were conducted under the scenario of administration of three bNAb infusions every eight weeks. For each model, the geometric mean PT_80_ was computed for 1,000 simulated participants, akin to those in the AMP trial, against panels of 31 Clade C AMP placebo viruses, 86 Clade C AMP VRC01 breakthrough viruses, and 68 Clade B AMP placebo viruses. For each simulated participant, the PT_80_ was calculated as the predicted steady-state serum concentration of 04_A06 divided by the geometric mean of the *in vitro* IC_80_ for the viruses in the AMP trial. These calculations assumed that 04_A06 displays the PK profile of either 3BNC117 or 10-1074^106^. The prevention efficacy was estimated by assuming that the PT_80_ needed for a specific level of prevention efficacy is twice as high as that found in non-human primate studies according to a meta-analysis, presenting a more conservative approach than that seen in the AMP trials^102,148^.

### Sequence annotation and clone assignment

For sequence analysis, chromatograms were first filtered based on a minimum Phred score of 28 and a length of at least 240 nucleotides (nt). Sequences were then processed using IgBLAST to annotate them, and trimmed to extract only the variable region, extending from FWR1 to the end of the J gene^132^. Base calls within the variable region with a Phred score below 16 were masked. Sequences containing over 15 masked nucleotides, stop codons, or frameshift mutations were excluded from subsequent analysis. Clonal analysis was conducted individually for each donor, where all productive heavy chain sequences were organized into groups based on identical VH/JH gene pairs and the pairwise Levenshtein distance of their CDRH3s was calculated. Clonal groups were formed starting from a randomly selected sequence, assigning sequences to a group if they had a minimum CDRH3 amino acid identity of 75% relative to the shortest CDRH3 in the group. 100 rounds of input sequence randomization and clonal assignment were performed and the outcome with the fewest unassigned (non-clonal) sequences was chosen for further analysis. The accuracy of all identified clones was cross-verified by the researchers, considering shared mutations and corresponding light chain information.

### Inference of the phylogenetic tree

The phylogenetic relationships among the sequences of clones 7, 9, and 1, as depicted in Fig. 2 were computed. To achieve this, nucleotide sequences within each clone were aligned using the multiple sequences alignment software Clustal Omega^149^, and the light chain sequence was concatenated with the heavy chain sequence to create an artificial, extended sequence. This approach allows tree reconstruction to incorporate information from both chains. Additionally, the germline sequence with a masked CDR3 region was included in the multiple sequence alignment. Phylogenetic tree inference was conducted using RaxML^150^, with the tree rooted at the germline sequence. Fig. 2d illustrates the three phylogenetic trees computed using the GTRGAMMA model. The segment lengths in the figure represent the number of mutations per nucleotide, with a reference length provided in the plot. The length of the first edge connecting the root (germline sequence) to the Most Recent Common Ancestor accounts for mutations only in the V and J genes of the heavy and light chains, as the CDR3 sequence is not available for the germline.

### Determination of intra- and interdonor bNAb similarities

The primary measure of similarity utilized in Fig. 2 is the fraction of common mutations from the germline in the VH gene. This similarity is calculated by aligning all V genes of the bNAbs from different donors using Clustal Omega^151^. The germline IGHV1-2 gene is also included in the alignment, enabling the determination of the number of mutations in each sequence relative to the germline. Notably, gaps are treated equivalently to other nucleotide characters when calculating mutations. For each pair of sequences, the number of common mutations relative to the germline is also determined. The similarity measure is then defined as follows:

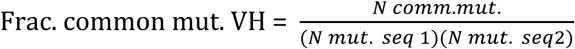

### Quantification and statistical analysis

Flow cytometry analyses and quantification were performed using FlowJo10 software. Statistical tests and analyses were done with GraphPad Prism (v7 and v8), Python (v3.6.8), R (v4.0.0) and Mircosoft Excel for Mac (v14.7.3 and 16.4.8). CDRH3 lengths, V gene usage and germline identity distributions for clonal sequences were assessed for all input sequences without further collapsing.

### Data availability

Nucleotide sequences of isolated VH1-2-encoded CD4bs bNAbs will be deposited at GenBank after review or acceptance of the manuscript. Further heavy and light chain sequences as well as NGS data of B cell repertoires from healthy individuals will be shared by the Lead Contact upon request. Cryo-EM maps and models have been deposited in the Electron Microscopy Data Bank (EMDB) and Protein Data Bank with accession codes: EMD-46649 and PDB ID 9D8V (unbound BG505), EMD-42363 and PDB ID 8ULR (05_B08-BG505v2), EMD-42364 and PDB ID 8ULS (01_D03-BG505), EMD-42365 and PDB ID 8ULT (04_A06-BG505), and EMD-42366 and PDB ID 8ULU (04_A06-PGDM1400-BG505). Coordinates for the 04_A06 Fab crystal structure have been deposited to the Protein Data Bank with accession code 8UKI. Aggregated clinical data are available upon request to the corresponding author (F.K.) provided that there is no reasonable risk of de-anonymizing study participants. Individual donor data cannot be shared due to privacy restrictions.

### Code availability

All computer code and data for the deep mutational scanning is publicly available on GitHub (https://github.com/dms-vep/HIV_Envelope_BF520_DMS_04-A06). Code implementation for *in silico* modelling of the predicted prevention efficacy is described in Gilbert et al.^152^ and publicly available at http://faculty.washington.edu/peterg/programs.html<u>?</u>.

## Acknowledgments

We thank all study participants for supporting our research by blood donation; members of the Klein, Bjorkman, and Bloom Labs for their support and inspiring discussions; Xualing Shi, Tongqing Zhou, and Linqi Zhang for sharing and providing the plasmids encoding for CD4bs-resistant pseudovirus strains; Anna Schmidt and Tina Bresser for lab management and assistance; We thank Songye Chen and the Caltech Cryo-EM Center and Jost Vielmetter and the Caltech Protein Expression Center for experimental support. A.T.D. was supported by an NSF Graduate Research Fellowship. We thank the Cologne Center for Genomics for sequencing and library preparation support; and the staff of the Animal Care Facility Weyertal at the University of Cologne. The panel of global HIV-1 pseudoviruses was obtained through the NIH AIDS Reagent Program. This work was funded by grants from the German Center of Infection Research (DZIF) to F.K and S.B., the German Research Foundation (DFG) CRC1279 and CRC1310, European Research Council (ERC) ERC-stG639961, and COVIM: NaFoUniMed-Covid19 to F. Klein. This work was also supported, in whole or in part, by the Bill & Melinda Gates Foundation (grant INV-002143) to F.K. and P.J.B. and (grant INV-036842) to M.S.S.. Under the grant conditions of the Foundation, a Creative Commons Attribution 4.0 Generic License has already been assigned to the Author Accepted Manuscript version that might arise from this submission. Research reported in this publication was also supported by the National Institute Of Allergy And Infectious Diseases of the National Institutes of Health under award number P01AI100148 and 1U54AI170856 to P.J.B., and R01AI140891 and U01AI169385 to J.D.B.. The content is solely the responsibility of the authors and does not necessarily represent the official views of the National Institutes of Health. P.L.M is supported by the South African Research Chairs Initiative of the Department of Science and Innovation and the National Research Foundation (Grant No 98341). This study was supported in part by the Bill & Melinda Gates Foundation’s Collaboration for AIDS Vaccine Discovery (CAVD; grant 1032144) to P.L.M.. This work was also supported by the European Research Council COG 724208 and ANR-19-CE45-0018 “RESPREP” from the Agence Nationale de la Recherche to A.W.. P.S. is supported by the Emmy Noether Programme of the German Research Foundation (DFG; project no. 495793173). BnAb 04_A06 has been exclusively licensed to Vir Biotechnology.

## Author Contributions

Conceptualization – L.G., A.D., F.K., P.J.B., H.B.G, C.K. and H. Gruell.; Methodology – L.G., C.K., H. Gruell., A.D., H. Gristick, C.R., J.D.B, P.F.S, A.M., A.W., and T.M.; Formal analysis – L.G., A.D., C.K., H. Gristick., A.P.W., C.R., N. D., M.S., P.F.S, A.M., A.W., L.Z. and T.M ; Investigation – L.G., A.D., F.K., P.J.B., H. Gristick, C.R., N.D., M.S., and J.D.B.; Resources – H. Gruell., P.F.S.; Writing original draft – L.G., A.D., H. Gristick, C.R., F.K., P.J.B., C.K., H. Gruell, A.M. and F.K.; Writing reviewing and editing – all authors; Supervision – F.K., P.J.B., H. Gristick, and H. Gruell.; Funding acquisition – F.K., P.J.B.

These authors contributed equally to this work: L.G. and A.D.

## Competing interest declaration

A patent application that comprises aspects of this work has been filed by the University of Cologne, listing L.G. and F.K. as inventors. H.G., P.S., and F.K. are listed as inventors on additional patent applications on HIV-1 neutralizing antibodies and have received payments from the University of Cologne for licensed patents. JDB and CER are inventors on Fred Hutch licensed patents related to viral deep mutational scanning. JDB consults for Apriori Bio.

## Materials & Correspondence

Further information and requests for resources and reagents should be directed to and will be fulfilled by the Lead Contact, Florian Klein.

## Inclusion & Ethics

Research has been conducted and authorships have been determined in alignment with the Global Code of Conduct for Research in Resource-Poor Settings.

## Supplementary information

**Supplementary Table 1: Demographical characteristics of analyzed HIV-1 elite neutralizers**

**Supplementary Table 2: Neutralizing activity of isolated mAbs against a HIV-1 pseudovirus screening panel**

**Supplementary Table 3: Isolated HIV-1-specific B cell clones and sequences from individual EN02**

**Supplementary Table 4: Neutralizing activity against the global pseudovirus panel and binding activity of isolated mAbs from EN02**

**Supplementary Table 5: Antiviral activity of 04_A06, 01_D03 and 05_B08 against the 119 multiclade pseudovirus panel**

**Supplementary Table 6: Antiviral activity of 04_A06, 01_D03 and 05_B08 against the 208 multiclade pseudovirus panel**

**Supplementary Table 7: Antiviral activity of 04_A06 against the Subtype C pseudovirus panel**

**Supplementary Table 8: Antiviral activity of 04_A06 and VRC07_523-LS_ against replication competent viral isolates**

**Supplementary Table 9: Cryo-EM data collection and refinements statistics**

**Supplementary Table 10: SGS-derived env sequences from monotherapy groups of *in vivo* experiments**

**Supplementary Table 11: SGS-derived env sequences from combination therapy groups of *in vivo* experiments**

**Supplementary Table 12: Neutralizing activity of 04_A06 against pseudoviruses generated from the AMP trials**

---

**Extended Data Fig. 1:**
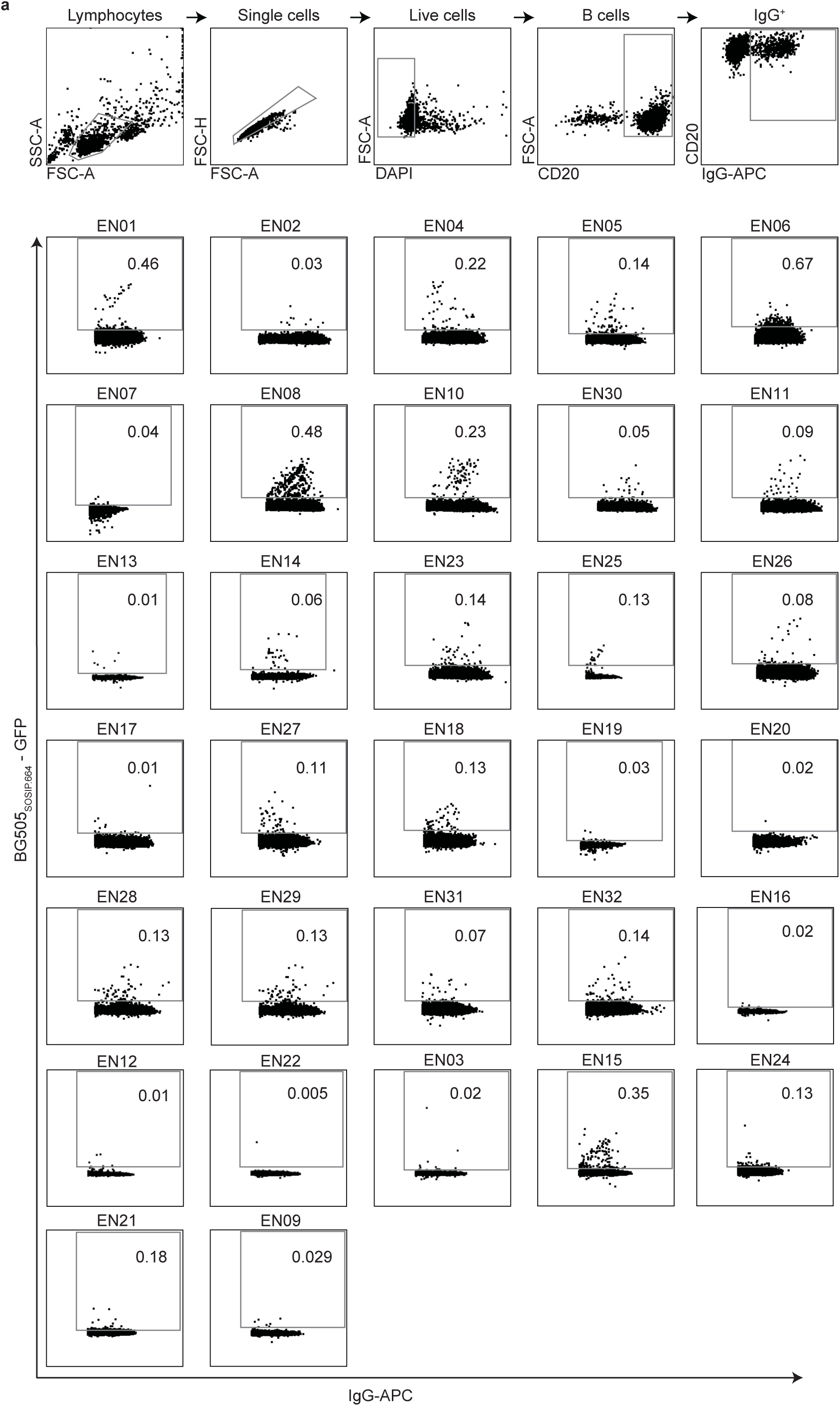
Gating strategy and single cell sorts of HIV-1-reactive B cell subsets. **a**, Individual FACS plots depicting sorting gates and frequencies of HIV-1-reactive, IgG^+^ B cells (in %) isolated from 32 donors. Numbers indicate the frequency of HIV-1 Env-reactive IgG+ B cells from the parental gate.

**Extended Data Fig. 2:**
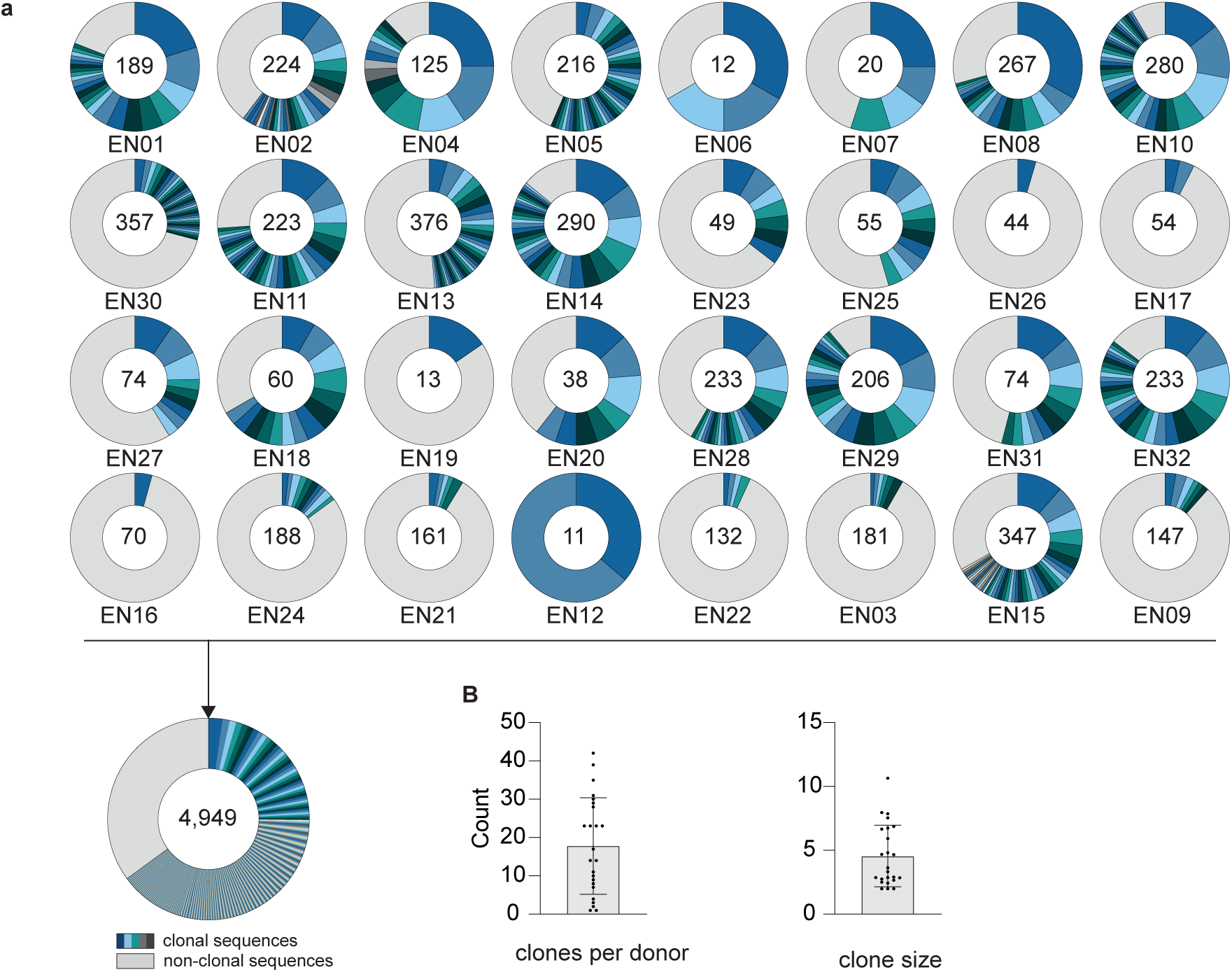
Analysis of isolated heavy chain sequences. **a**, Clonal relationship of heavy chain sequences amplified from single HIV-1-reactive IgG+ B cells isolated from 32 donors. Individual clones are colored in shades of blue, gray, and white. In the center of each pie chart, numbers of productive heavy chain sequences are illustrated. Presentation of clone sizes are proportional to the total number of productive heavy chain sequences per clone. **b,** Dot plot bar graphs display the mean ± SD.

**Extended Data Fig. 3:**
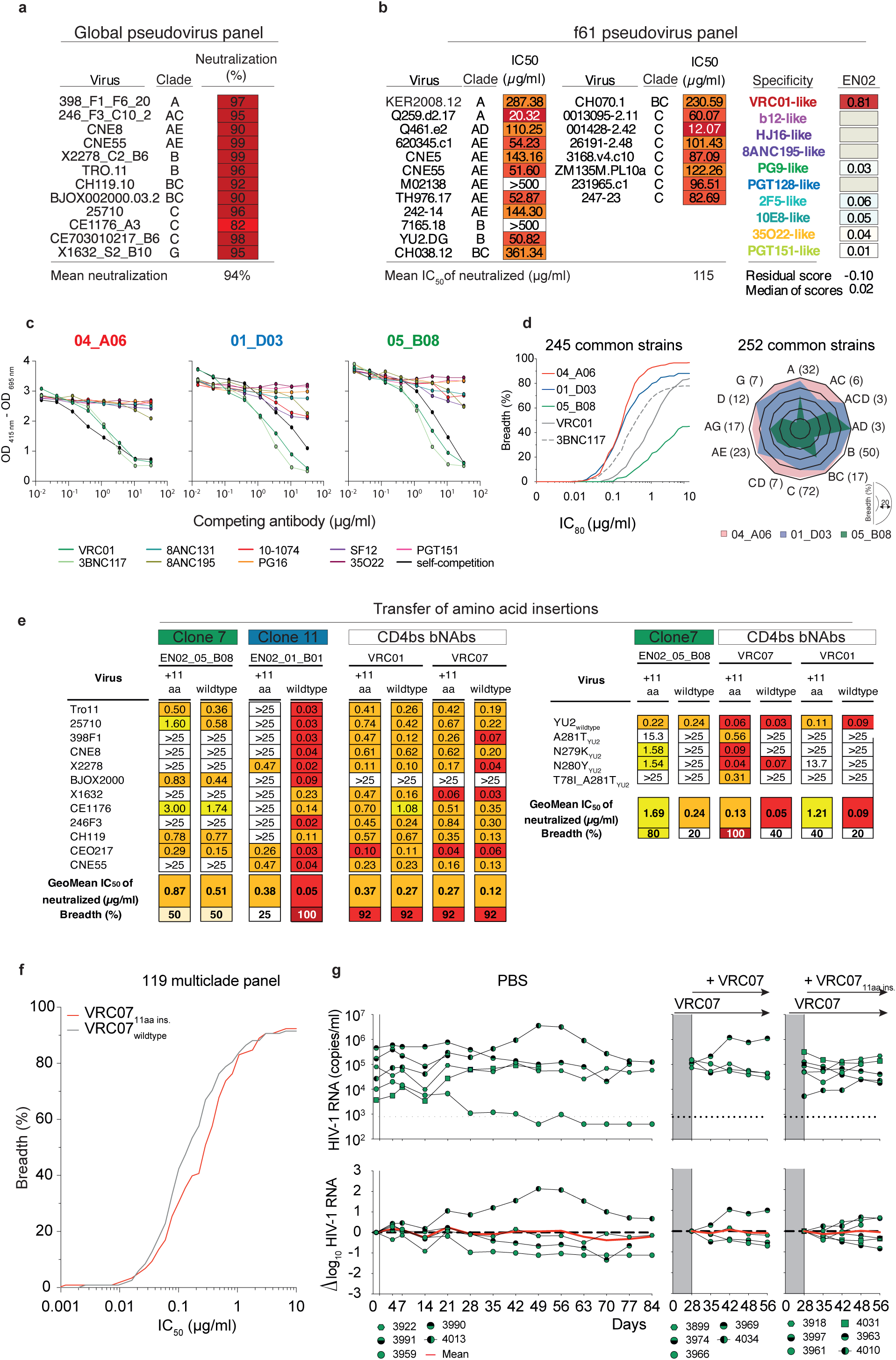
Neutralization and binding profile of serum, isolated and chimeric mAbs from donor EN02. **a**, Neutralizing activity of EN02 serum IgG against the HIV-1 global and **b**, f61 pseudovirus panel retrieved from^65^. Right panels show delineation scores of f61 panel-based computational epitope mapping. **c**, Interference of indicated bNAbs with reference bNAbs targeting known epitopes on the HIV-1 Env trimer measured by competition ELISAs. **d**, Neutralization breadth (%) and potency (GeoMean IC_80_) of representative bNAb 04_A06 (clone 9; red), 01_D03 (clone 11, blue); and 05_B08 (clone 7, green) against ≥245 pseudovirus strains. Samples were tested in duplicates. Neutralization data for reference bNAbs was retrieved from CATNAP database^128^. **e**, The elongated heavy chain FWRH1 of 04_A06 was engrafted into clonally distinct bNAbs isolated from donor EN02 and CD4bs reference bNAbs. The antiviral activity of wildtype and chimeric versions of isolated and reference mAbs was determined against the global HIV-1 panel and common CD4bs escape pseudovirus variants. **f**, Breadth (%) and potency (IC_50_) of the chimeric VRC07 antibody version against the 119 multiclade panel. **g**, Sequential treatment with the chimeric VRC07 antibody version (VRC07_11aa ins_) in HIV-1_YU2_-infected humanized mice following viral rebound during VRC07 monotherapy. This approach included maintaining VRC07 monotherapy while integrating VRC07_11aa ins._ in the treatment regimen. Mice treated with a double dose of VRC07 (1mg) were included as a control group. Dashed lines in the top graphs indicate the lower limit of quantitation of the qPCR assay (LLQ) (784 copies/ml). Red lines display the average log_10_ changes compared to baseline viral loads (day 0).

**Extended Data Fig. 4:**
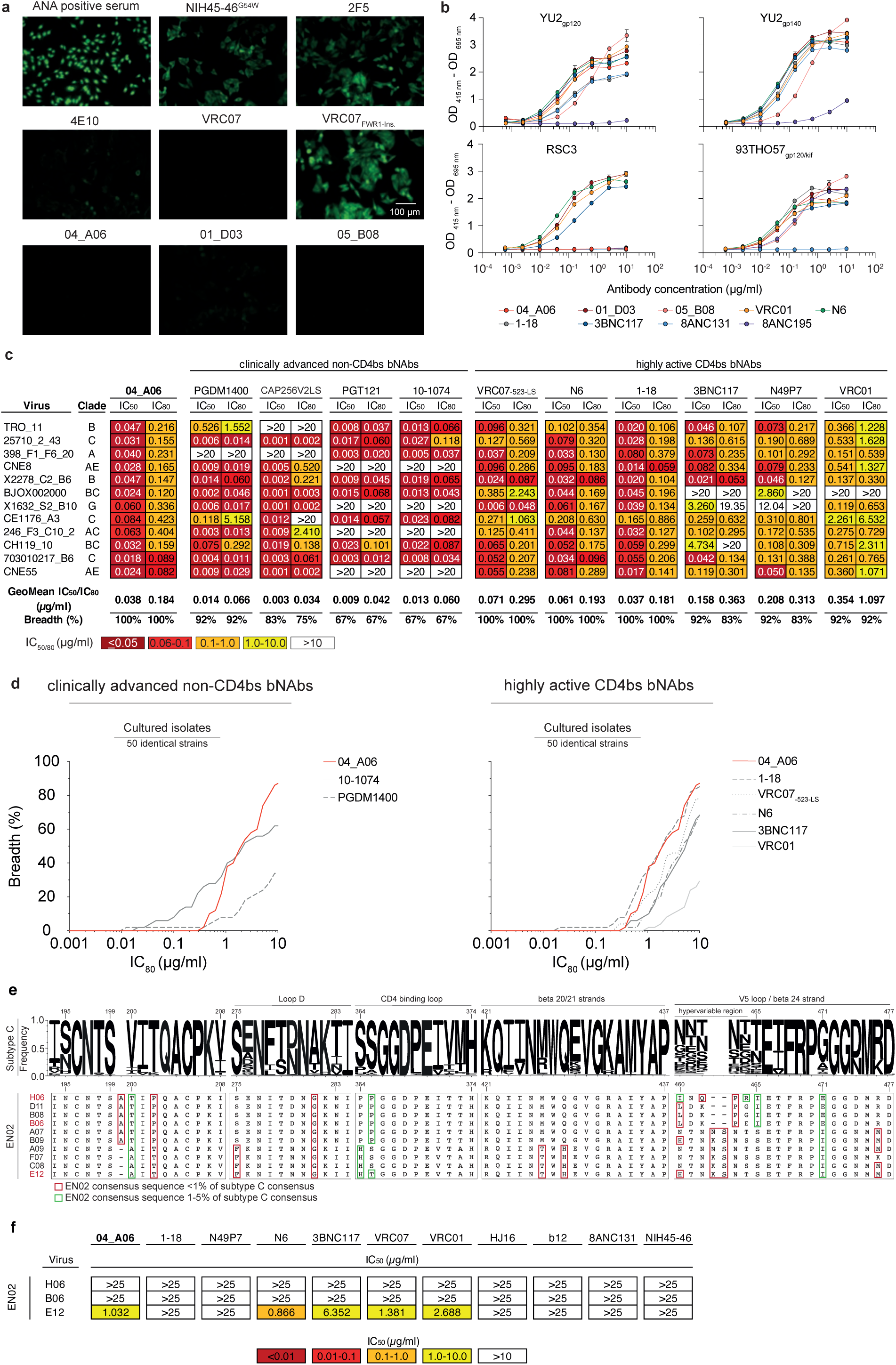
Autoreactivity and antiviral activity of isolated bNAbs from donor EN02. **a**, Reactivity of indicated antibodies against HEp-2 cells. Antibodies were tested at a concentration of 100 µg/ml. **b**, Binding profiles of isolated bNAbs from donor EN02 against indicated HIV-1 Env antigens. **c**, Neutralization activity of bNAb 04_A06 and reference bNAbs against the HIV-1 global pseudovirus panel. Samples were tested in duplicates. Neutralization data of reference bNAbs were retrieved from CATNAP database^128^. **d**, Neutralization coverage (%) and potency (IC_80_) of 04_A06 against a panel of 50 donor-derived bulk outgrowth isolates. Neutralization data of reference bNAbs against replication competent viruses were retrieved from^36^. Samples were tested in duplicates. **e,** Amino acid frequency at selected sites across 744 clade C sequences from the Los Alamos National Laboratory (LANL) HIV database is shown (top; letter height reflects frequency). The lower panels depict corresponding positions in plasma-derived single genome sequencing (SGS) *env* sequences from donor EN02. Red and green boxes highlight amino acid residues present in <1% and 1–5%, respectively, of LANL clade C sequences. Numbering corresponds to HIV-1 HXB2 reference strain. **f,** Neutralization sensitivity of selected pseudoviruses carrying EN02 *env* sequences shown in **e**.

**Extended Data Fig. 5:**
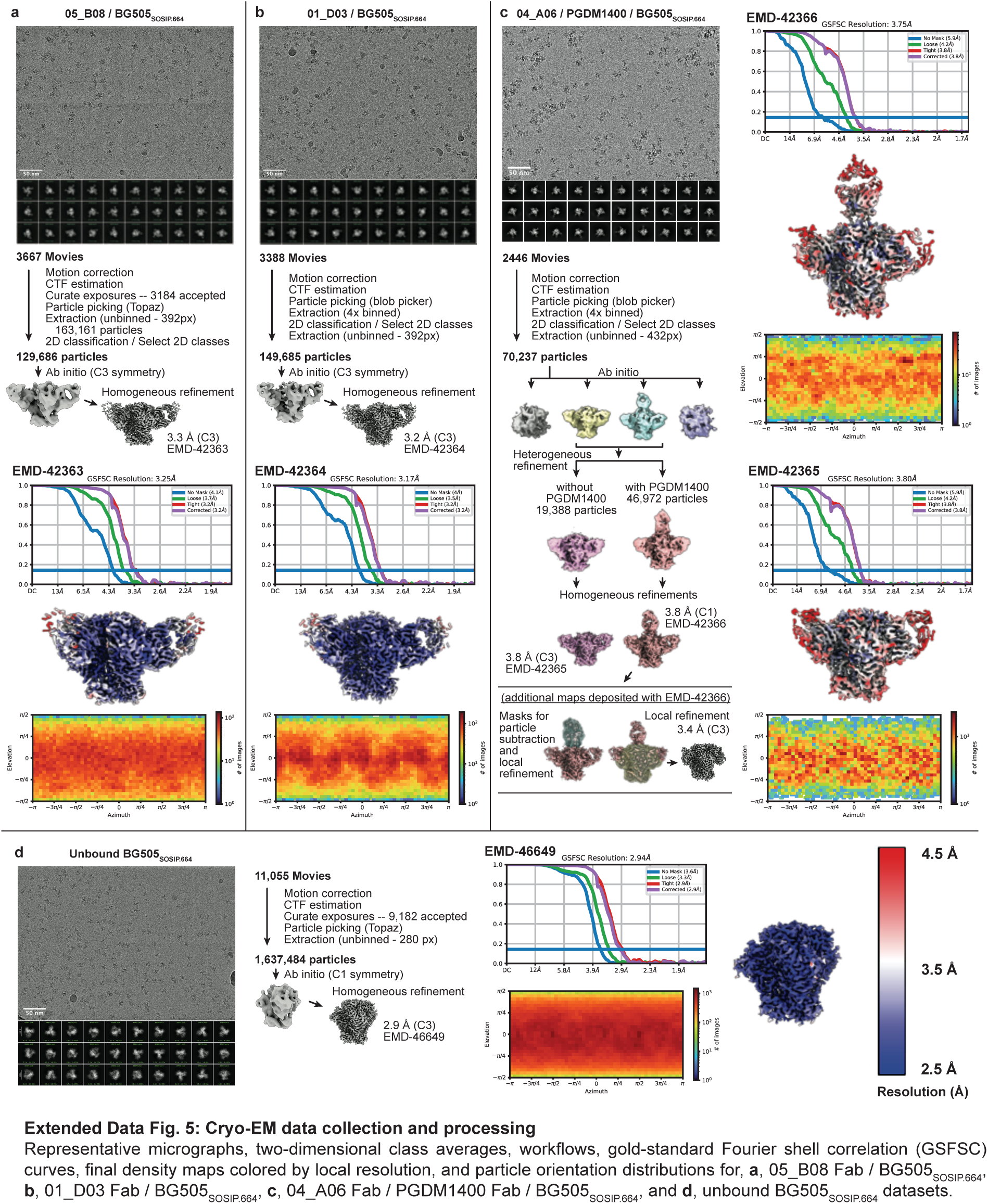
Cryo-EM data collection and processing. Representative micrographs, two-dimensional class averages, workflows, gold-standard Fourier shell correlation (GSFSC) curves, final density maps colored by local resolution, and particle orientation distributions for **a**, 05_B08 Fab / BG505_SOSIP.664_, **b**, 01_D03 Fab / BG505_SOSIP.664_, **c**, 04_A06 Fab / PGDM1400 Fab / BG505_SOSIP.664_, and **d**, unbound BG505_SOSIP.664_ datasets.

**Extended Data Fig. 6:**
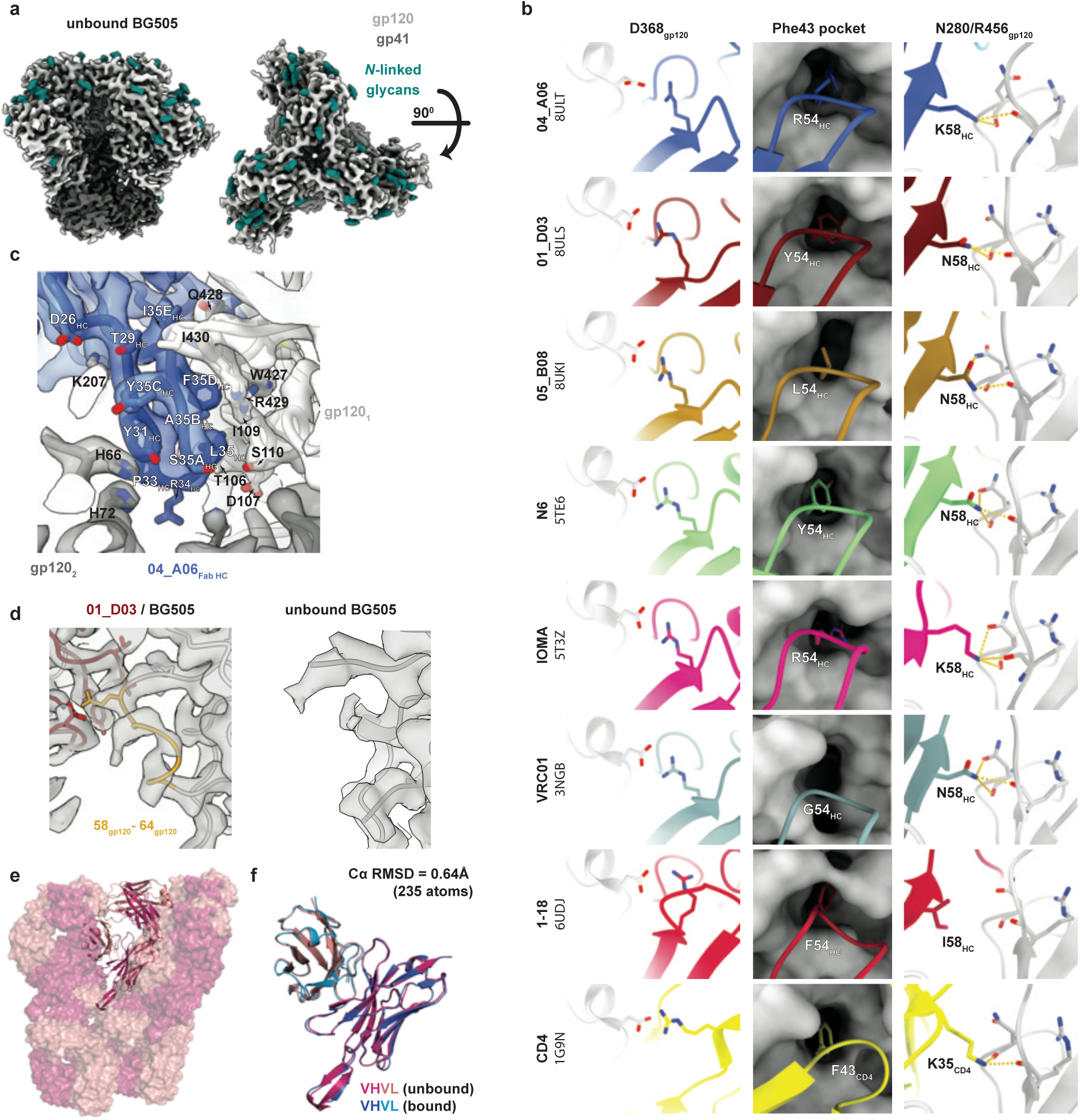
Structural analysis of CD4bs antibodies from EN02 with other CD4bs antibodies and unbound BG505. **a**, EM density of unbound BG505_SOSIP.664_ structure showing side and top views. **b**, Canonical interactions of CD4 and CD4bs bNAbs with Env: D368_gp120_, Phe43 pocket, and N280_gp120_ / R456_gp120_. **c**, Close up view of 04_A06’s CDRH1 interactions (blue) with the primary gp120 protomer (gp120_1_). **d**, EM density for gp120 residues 58-64 on the adjacent protomer (gp120_2_) as observed in complex with 01_D03 Fab compared to unbound BG505_SOSIP.664_. **e**, Crystal packing environment of 04_A06 Fab. **f**, Overlay of V_H_-V_L_ domains in bound (from Fab-SOSIP cryo-EM structure) and unbound (from Fab crystal structure) structures.

**Extended Data Fig. 7:**
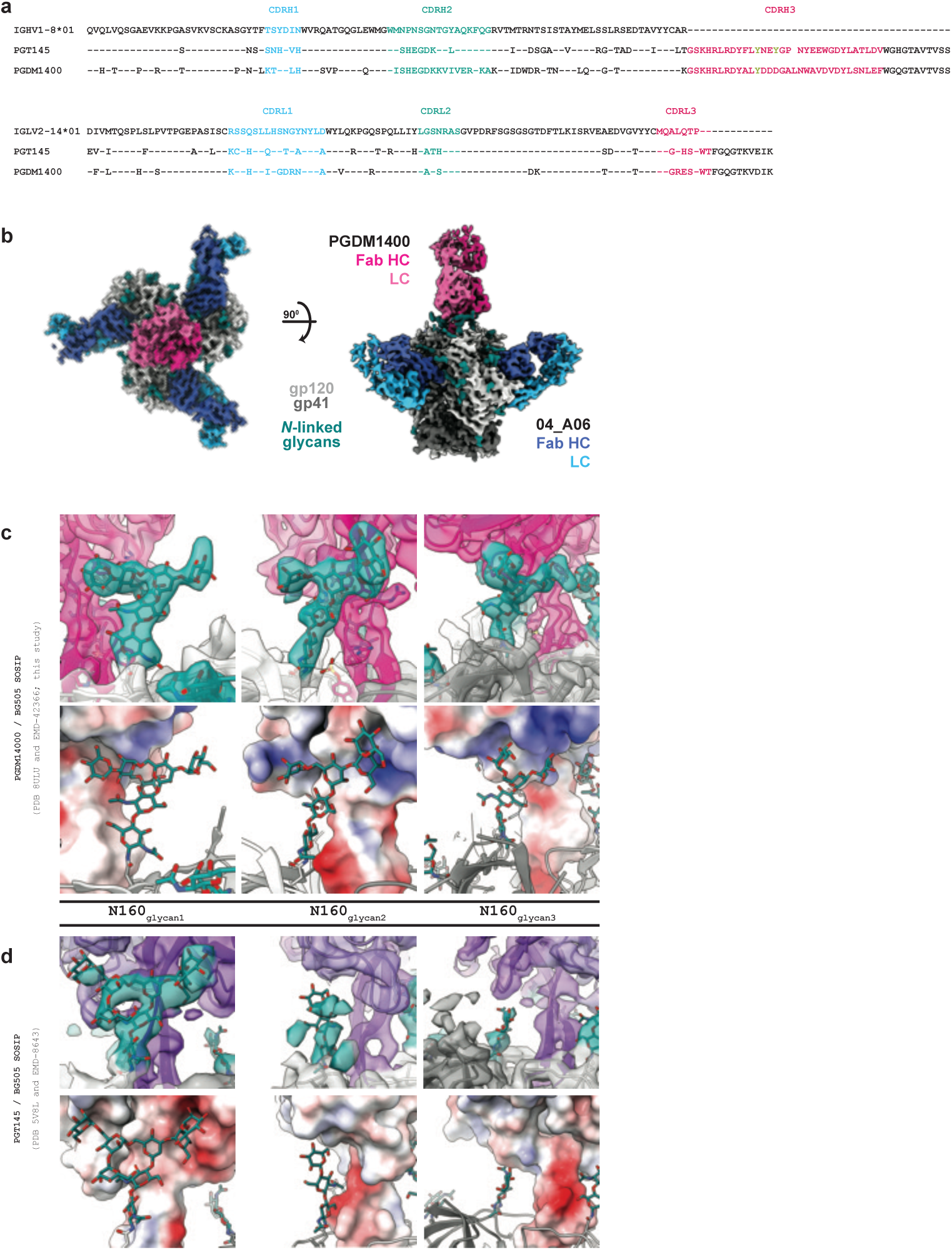
Structural analyses of PGDM1400. **a**, Sequence alignment of heavy and light chain V genes from bNAbs PGT145 and PGDM1400. Sulfated tyrosine residues located in the CDRH3 are indicated by a green “Y”. **b**, Structure overview showing EM density of BG505_SOSIP.664_ in complex with 04_A06 and PGDM1400 Fabs. EM density for N160 glycans (top) and a Fab electrostatic surface calculation (bottom) for PGDM1400 (**c**) and PGT145 (**d**). Glycans labeled as glycan 1,2, and 3 correspond to the N160_gp120_ glycans from each of the three Env protomers^82^.

**Extended Data Fig. 8:**
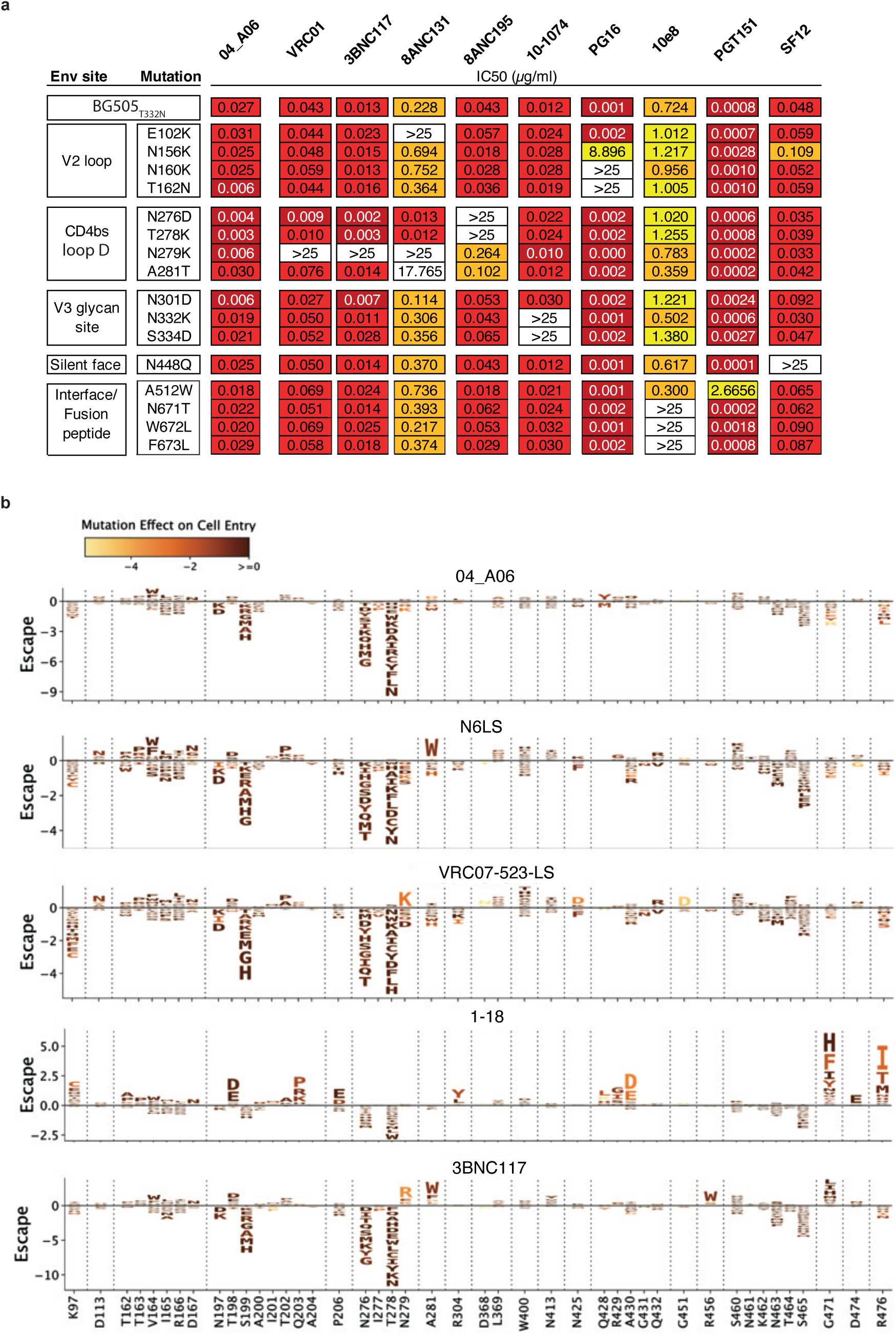
Restriction of viral escape *in vitro*. **a**, Neutralizing activity of 04_A06 against BG505_T332N_ and common escape pseudovirus mutants. The panels indicate the IC_50_ of 04_A06 against the respective pseudovirus variant. **b**, Logo plots illustrating the extent of neutralization escape caused by mutations in the HIV-1 Env strain BF520 for antibodies 04_A06, N6-LS, VRC07_-523-LS_, 1-18 and 3BNC117. The heights of individual letters represent the effect of that amino-acid mutation on antibody neutralization, with positive heights (letters above the zero line) indicating mutations that cause escape, and negative heights (letters below the zero line) indicating mutations that increase neutralization. Letters are colored by the effect of that mutation on pseudovirus cell entry, with yellow corresponding to reduced cell entry and brown corresponding to neutral effects on cell entry. The y-axis scales for each antibody are independent. Only key sites are illustrated. See https://dms-vep.org/HIV_Envelope_BF520_DMS_04-A06/htmls/04-A06_mut_effect.html, https://dms-vep.org/HIV_Envelope_BF520_DMS_04-A06/htmls/N6_LS_mut_effect.html, https://dms-vep.org/HIV_Envelope_BF520_DMS_04-A06/htmls/VRC07_523_LS_mut_effect.html, https://dms-vep.org/HIV_Envelope_BF520_DMS_04-A06/htmls/1-18_mut_effect.html, and https://dms-vep.org/HIV_Envelope_BF520_DMS_04-A06/htmls/3BNC117_mut_effect.html for interactive versions of the escape maps.

**Extended Data Fig. 9:**
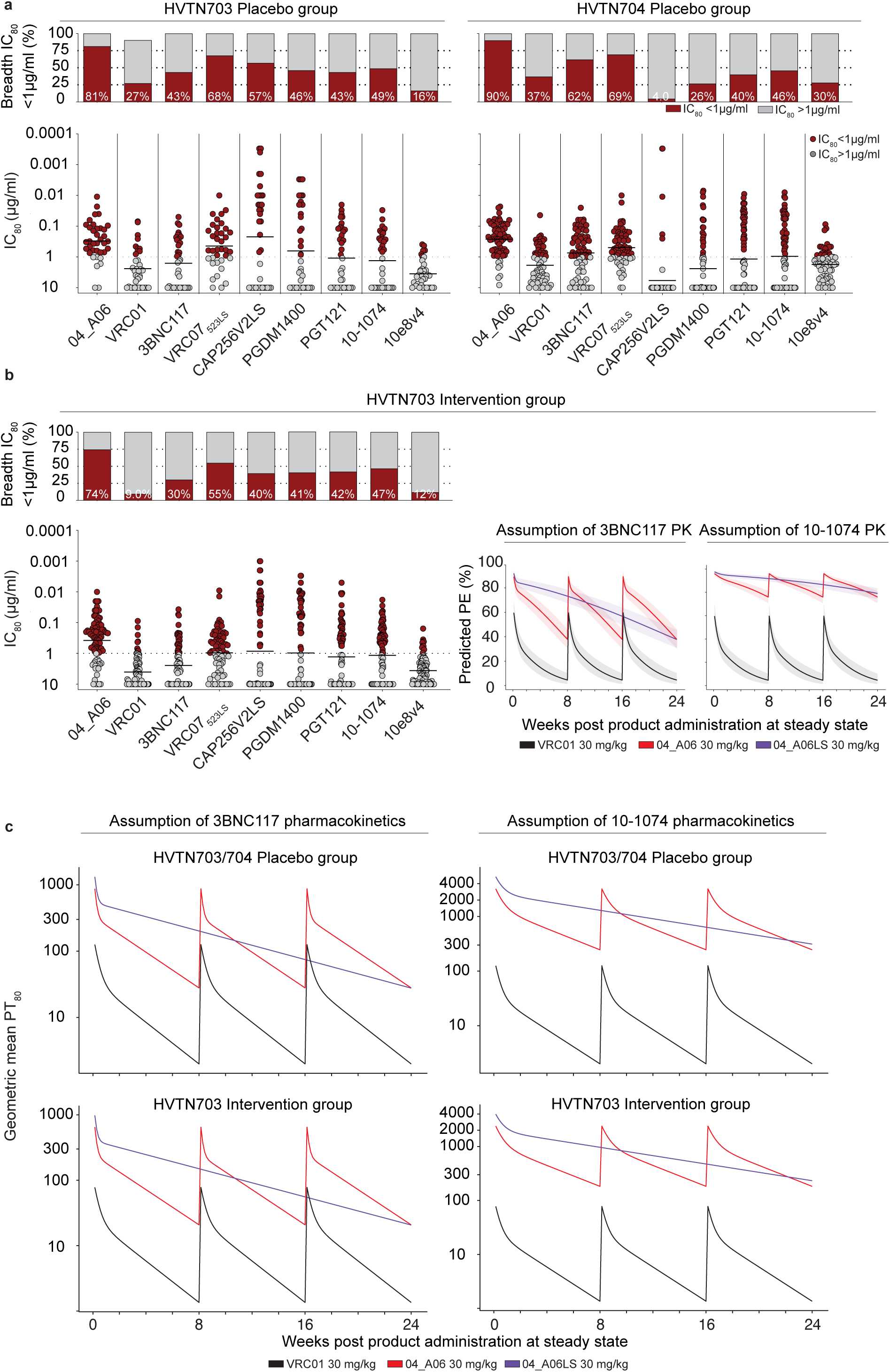
Neutralization profile and predicted PE against AMP trial pseudoviruses. Antiviral activity of 04_A06 and reference bNAbs against pseudoviruses generated from **a**, the HVTN703 and HVTN704 placebo group and **b**, HVTN703 intervention group, respectively. Bar graphs display the breadth (%) at the established threshold of protection (IC_80_ ≤ 1 µg/ml). Dot plots illustrate the potency of tested bNAbs (IC_80_) against each pseudovirus strain. Predicted HIV-1 PE of 04_A06 over time against HVTN703 intervention group pseudoviruses in comparison to VRC01. Left graph illustrates the PE of 04_A06 (red) and 04_A06LS (purple) under the assumption of 3BNC117 pharmacokinetics. The right graph displays the PE of 04_A06 and 04_A06LS under the assumption of 10-1074 pharmacokinetics. **c**, Illustration of the geometric mean PT_80_ against placebo and intervention group viruses. The geometric mean PT_80_ for each time point was determined by dividing the geometric mean of the predicted serum concentration of bNAbs in recipients at each time point during steady state (as simulated via PK modeling for each bNAb, as outlined in the methods section) by the geometric mean IC_80_ of the bNAb against viruses present in the specified AMP trial. Calculations were performed over time after three 8-weekly infusions of 04_A06 at 30 mg/kg or a single infusion of 04_A06LS at 30 mg/kg. In the scenario involving the LS-modified version, predictions were made under the premise that 04_A06LS has a 2.5-fold higher half-life than 04_A06. Black lines indicate the PT_80_ for VRC01. Data for reference bNAbs were retrieved from CATNAP database^127^ and^112^.

**Extended Data Fig. 10:**
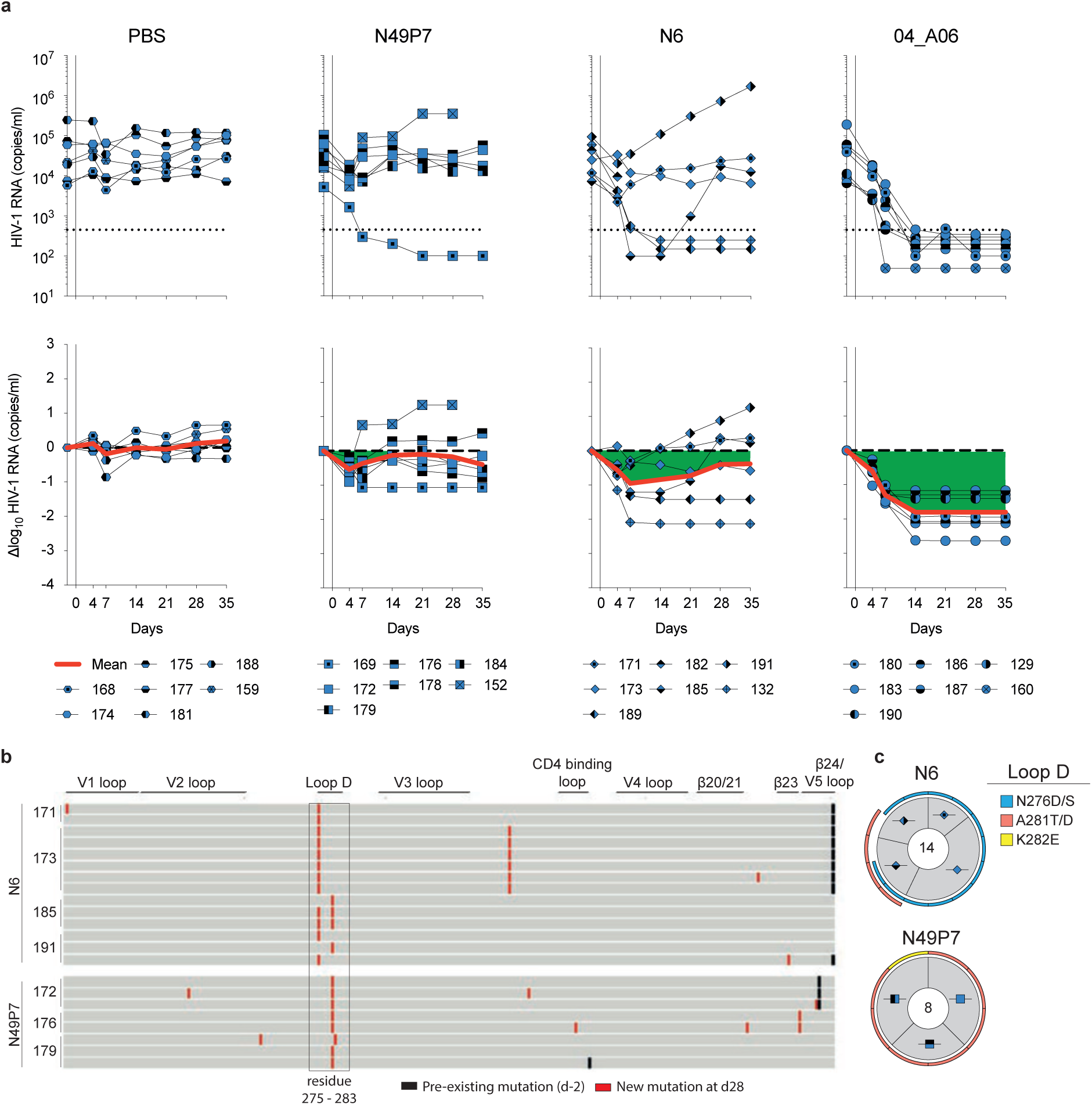
***In vivo* antiviral activity of 04_A06 in comparison to highly-active CD4bs bNAbs a,** Investigation of the antiviral activity of N49P7, N6, and 04_A06 monotherapy in HIV-1_YU2_-infected humanized mice (NXG-HIS). Graphs illustrate the absolute HIV-1 RNA plasma copies/ml (top) and relative log_10_ changes from baseline viral loads (bottom) after initiation of bNAb therapy. Dashed lines (top graphs) indicate the lower limit of quantitation of the qPCR assay (LLQ) (451 copies/ml). Red lines display the average log_10_ changes compared to baseline viral loads (day −2). **b,** Alignment of plasma SGS-derived *env* sequences identified from day −1 (black bars) and after viral rebound from day 28 (red bars). **c,** Analyses of single HIV-1 plasma *env* sequences from HIV-1_YU2_-infected humanized mice obtained on day 28 after bNAb treatment initiation. Total number of analyzed sequences is indicated in the center of each pie chart. Mice are labeled according to icon legends in a. Colored bars on the outside of the pie charts indicate mutations in Loop D.

